# Bats Could Learn Three-Dimensional Echolocation Using Echoacoustic Flow

**DOI:** 10.64898/2026.10.22.760884

**Authors:** Jan Steckel

**Affiliations:** Cosys-Lab, Faculty of Applied Engineering, University of Antwerp, Antwerp, Belgium; Flanders Make Strategic Research Centre, Lommel, Belgium

## Abstract

Echolocating bats can find prey and avoid obstacles in three dimensions using sound alone. They infer the direction of an echo from the way the emitter, the head and the outer ears filter sound, together described by the echolocation-related transfer function (ERTF) [1–3]. This filter differs between species and is likely to change as the ears grow [2, 4], so it has to be calibrated by experience. Barn owls and humans recalibrate auditory space after their ears are altered, instructed by vision in the owl [5, 6] and by a so far unidentified teacher in humans [7, 8]. How a bat flying in the dark could calibrate its ERTF is unknown [9]. Here we show that a simulated bat-like agent can learn its ERTF in azimuth and elevation from its echoes and movements alone, using echoacoustic flow, without labeled echo directions or an external teacher such as vision. The agent encodes each call into a structured latent scene, moves it with a physics-based geometric warp and predicts the next echoes through a differentiable renderer. It learns its ERTF efficiently and relearns it quickly after its outer ears change. The results form a first functional model of how bats could learn spatial hearing and yield testable hypotheses for behavioral experiments.

## 1 Introduction

Echolocating bats perceive their surroundings by listening to and analyzing the echoes of their own calls, and they do so in three dimensions, often in complete darkness [10, 11]. The range of a reflector is easy to infer from the delay of its echo, but its direction has to be read from how the echo differs between the two ears and across frequency. The head, the outer ears, the tragus and, in many species, the noseleaf together act as a direction- and frequency-dependent filter. On the receiving side, this filter is called the head-related transfer function (HRTF) [1–3]. Combined with the spatiospectral directivity pattern of the emitter, it forms what we have called the echolocation-related transfer function (ERTF), (or active HRTF [12]). Both the level differences between the ears and the frequency spectra received by each ear vary with azimuth and elevation [1]. Both azimuth and elevation can be inferred from the received echo spectra, and elevation in particular is inferred from spectral notches shaped by the pinna and the tragus [13, 14]. Deflecting the tragus impairs judgments of the vertical position of targets much more than of their horizontal position [15, 16].

To use these cues for 3D localization, a bat needs to know its own ERTF, and this knowledge has to be calibrated by experience. Ear shapes differ strongly between species [2] and individuals. Indeed, in humans the transfer functions of the outer ears differ significantly between individuals [17] (detailed within-species data for bats are lacking). The auditory periphery of juvenile bats matures over the first weeks of life, possibly in step with the growth of their outer ears [4]. The mapping from spectral cues to directions therefore has to be learned, and relearned whenever the ears change. Other animals show that such recalibration is possible: young barn owls with one ear plugged adjust their sound localization during a sensitive period [18], and owls raised with displacing optical prisms shift their auditory space map to match their vision [5]. Humans fitted with ear molds regain their ability to localize sounds in elevation over a period of weeks [7, 19], and ferrets with one ear plugged relearn to localize at a rate set by the amount of training [20]. In the owl, vision is the instructive signal [6]. Ferrets, however, also relearn without vision [20], humans relearn for directions outside their visual field [21], and what serves as the teacher remains an open question [8]. Many bats hunt and navigate in darkness, where vision cannot serve as the teacher, although they do use vision when it is available [22].

We hypothesize that a candidate teacher is the bat’s own movement. When a sensor moves, its received exteroceptive signals change in a way that is determined by the sensor’s motion and by the layout of the world, and this lawful change can in principle reveal properties of the sensor itself [23, 24]. Aytekin and colleagues proposed exactly this mechanism for learning sound localization: a naive organism could learn auditory space from the relation between its acoustic inputs and its motor actions [9]. Robots have learned auditory-motor maps online without a visual teacher [25], a crude, innate sense of left and right turns out to be enough to bootstrap a spatial map that includes elevation [26], and a model of a sonar agent has learned to localize targets with active head rolls, without labeled target directions [27]. These studies, however, consider sounds from external sources, or they learn a mapping from cues to directions or a localization policy, rather than an explicit transfer function of the ears. In echolocation, the situation is more favorable, as the bat can act as its own sound source. As it flies through a static scene, the echoes from the surrounding reflectors shift in range and direction in a predictable way, a pattern known as echo-acoustic flow [28, 29]. Bats use this flow to control their flight [30, 31], it reshapes the range maps in their auditory cortex [32], and bats combine the echoes of successive calls to predict where a moving target will be [33]. In robotics, acoustic flow has been used extensively to steer mobile platforms [34, 35] and, recently, to calibrate the unknown directivity of a man-made sonar array sensor from its own ego-motion [36]. Learning a sensor model by predicting the sensory consequences of one’s own movement has deep roots in neuroscience [37, 38], and in computer vision it allows depth to be learned from video without any labels [39].

Beyond a teacher, we think that a biologically plausible solution for ERTF learning should meet three conditions. First, it should be able to learn online. Learning from a short memory of recent calls asks much less of an animal than storing and reprocessing a long history of sensory data, which is costly in neural tissue [40], especially because auditory sensory memory has been shown to last only a few seconds [41]. Second, the agent’s motion has to be rich enough to disambiguate the various dimensions relevant to the learning task (in our case: range, azimuth and elevation). For example, motion within the horizontal plane leaves the up/down sign of elevation ambiguous, and rotations out of that plane can resolve it. Bats are observed to make such rotations: they move their heads and ears while tracking targets [42], and in flight they rotate their bodies out of the horizontal plane [43] duing banking. In a model of echolocation, head rolls improve localization in elevation [27]. Third, the animal has to survive the learning period, which requires an operational sensor. Obstacles, however, can be avoided with simple cues that require no explicit localization and hence no detailed knowledge of the ERTF, such as the interaural level difference of the first echo [44, 45].

In this paper, we show that these aforementioned ingredients suffice to learn the ERTF in azimuth and elevation. We simulate a bat-like agent with one emitter, two ears, the numerically simulated ERTF of *Phyllostomus discolor* [3], a frequency-modulated call and a cochlear front end. From each call, a learned encoder infers the distribution of echo energy over range and direction, which is subsequently warped using the self-motion of the agent. This warped representation is then rendered using a learned ERTF into the cochleograms the agent expects to hear in the near future. The mismatch with the echoes it actually hears is the only training signal in this process, and no ground truth about the distribution of the reflectors enters the learning. Starting from a weak prior that knows only that the left ear favors the left side and the right ear the right side, the agent learns its ERTF in azimuth and elevation, online, from a short replay memory of recent calls. Its single-call scene estimates also encode where the reflectors around it are in three dimensions. In ablation experiments we show that when the self-motion readings are taken from other moments of the drive, the learning of the ERTF fails. After we rotate the ears, the agent relearns the new ERTF within about 1,000 calls, whereas the first learning needed 26,000 to 32,000 calls to improve by the same amount.

The results show that, in static scenes, the prediction of self-generated echoes under acoustic flow suffices to learn the ERTF. The model thus offers a first-order account of how bats could learn their ERTF without an external teacher, and it leads to predictions that can be tested in bats and on sonar robots.

### 2 Learning ERTFs by Predicting Echoes

The simulated agent in this paper is a ground robot with a bat-like sonar head on a pan-tilt-roll unit (Fig. 1a). It moves along smooth paths through worlds of point reflectors scattered between the floor and a height of 3 m (Supplementary Fig. 2), emits a frequency-modulated call every 10 cm of travel, and hears the echoes through two ears whose directivity, together with that of the emitter, is the simulated ERTF of *P. discolor* [3]. A cochlear filterbank turns the echoes of each call into a binaural cochleogram, the echo energy per frequency band and range. Apart from these cochleograms, the only data the agent receives are readings of how it moved between calls: the displacement measured by its wheels, the rotation measured by a gyroscope and the angles of its head joints, all with specified observation noise. The robot never receives true positions, spatial maps of the reflectors or its true ERTF.

**Figure 1:**
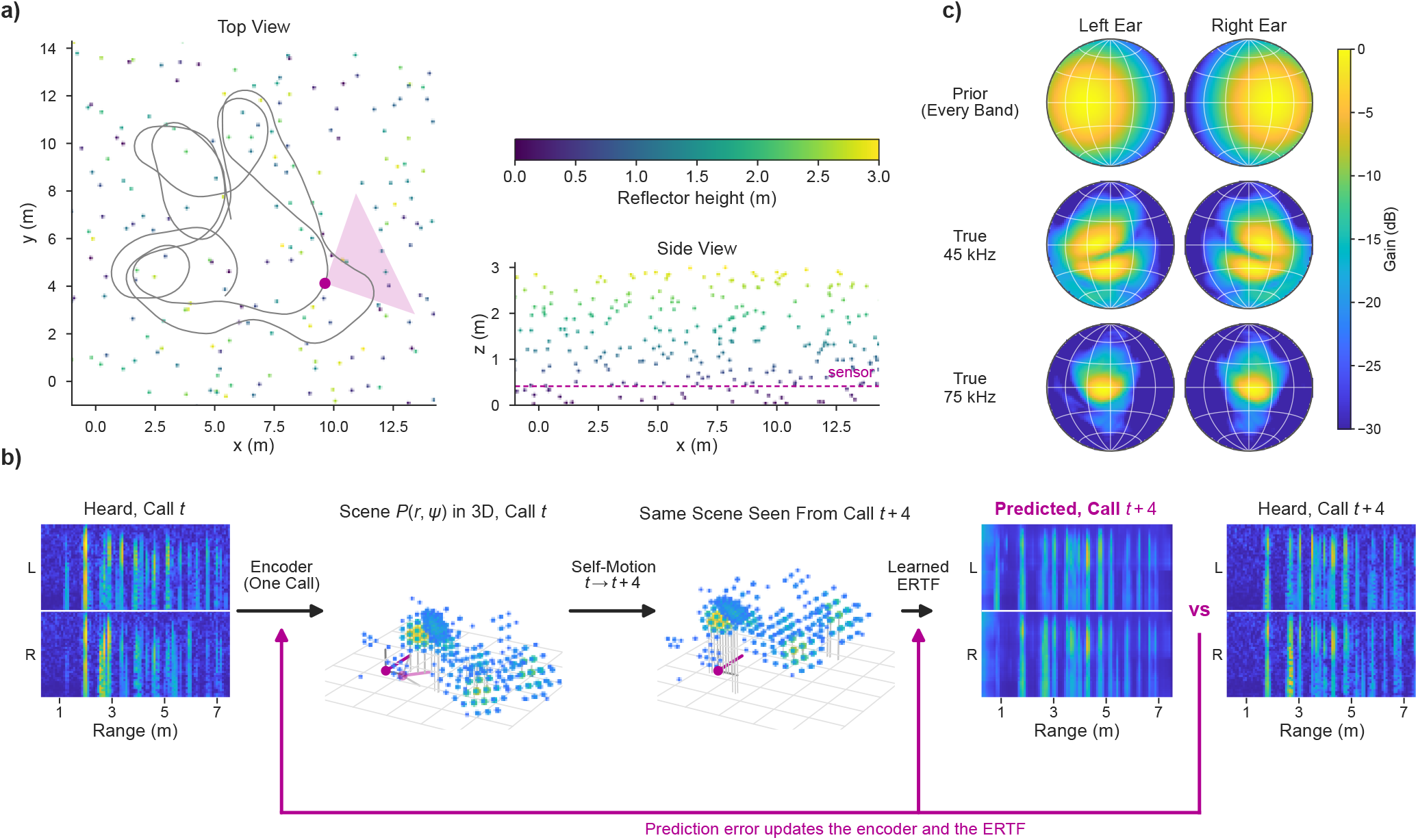
Learning the Ears by Predicting Echoes. **a**, One of the unseen validation worlds from above and from the side: point reflectors (color: height), the smooth path of the agent (gray) and its sensor with the field of view at the call shown in b (magenta). **b**, One learning step, with the model’s actual intermediate results for that call. The encoder turns the binaural cochleogram of call *t* into an EnergyScape *P* (*r, ψ*), a three-dimensional scene around the sensor: every point is a cell at range *r* and direction *ψ*, sized and colored by its energy (stems: height above a reference plane, magenta axes: the sensor, pale: its pose at call *t* + 4). The measured self-motion moves this scene rigidly into the frame of call *t* + 4, where the learned ERTF renders it into a predicted cochleogram. The error with respect to the cochleogram that is actually heard updates the encoder and the ERTF together. **c**, The weak prior the agent starts from, the same in every band, next to the true ERTF it has to find at 45 and 75 kHz (both ears, frontal hemisphere seen from behind the head). The true ERTF and the reflector positions are shown for orientation only and are never used for learning.

The learning is driven by the requirement that what the agent infers from one call predicts what it hears over the next few calls (Fig. 1b). From a single cochleogram, a learned encoder infers an EnergyScape *P*(*r, ψ*), the echo energy as a function of range *r* and direction *ψ* in the coordinate frame of the head. Because the reflectors are assumed to not move between calls, the measured self-motion fixes where every part of this scene must appear at each of the next calls. This transformation, the called the warping opeation, has no free parameters: it is an exact rigid transform of the scene, evaluated on the grid by interpolation, using the differential equations of acoustic flow [34]. A renderer with prescribed geometric spreading, learned absorption and range blur, and a learned representation of the transfer function then predicts the cochleograms of those calls, and the error between the prediction and the observed cochleogram is used as the training signal. As the emitter and ear directivities always act together on an echo, what the agent can learn is their product, the ERTF, per cochlear band [46]. About its ears, the agent knows at the start only that its left ear favors the left side and its right ear the right side: a smooth lobe per ear, the same in every band, without spectral detail and without any information on elevation (Fig. 1c), called the prior ERTF in what follows.

## 3 The ERTF Is Learned While Driving

We first let the agent learn online, similar to real-world animal learning: it drives through the 40 training worlds subsequently, takes one learning step after every call and keeps only a short replay memory: the training windows (17 consecutive calls each) that end within its last 65 calls, or 81 calls of raw history (8 m of travel, see Methods for details). Fig. 2a shows the left ear and the interaural level difference (ILD) at fixed moments of this drive. After 1,000 calls, the two broad lobes of the prior have contracted toward the correct directions. After a few thousand, the main lobes and the first spectral notches appear, and at the end, the learned ERTF reproduces the structure of the true one, including the asymmetry between up and down that the prior ERTF did not contain (Supplementary Video 1).

**Figure 2:**
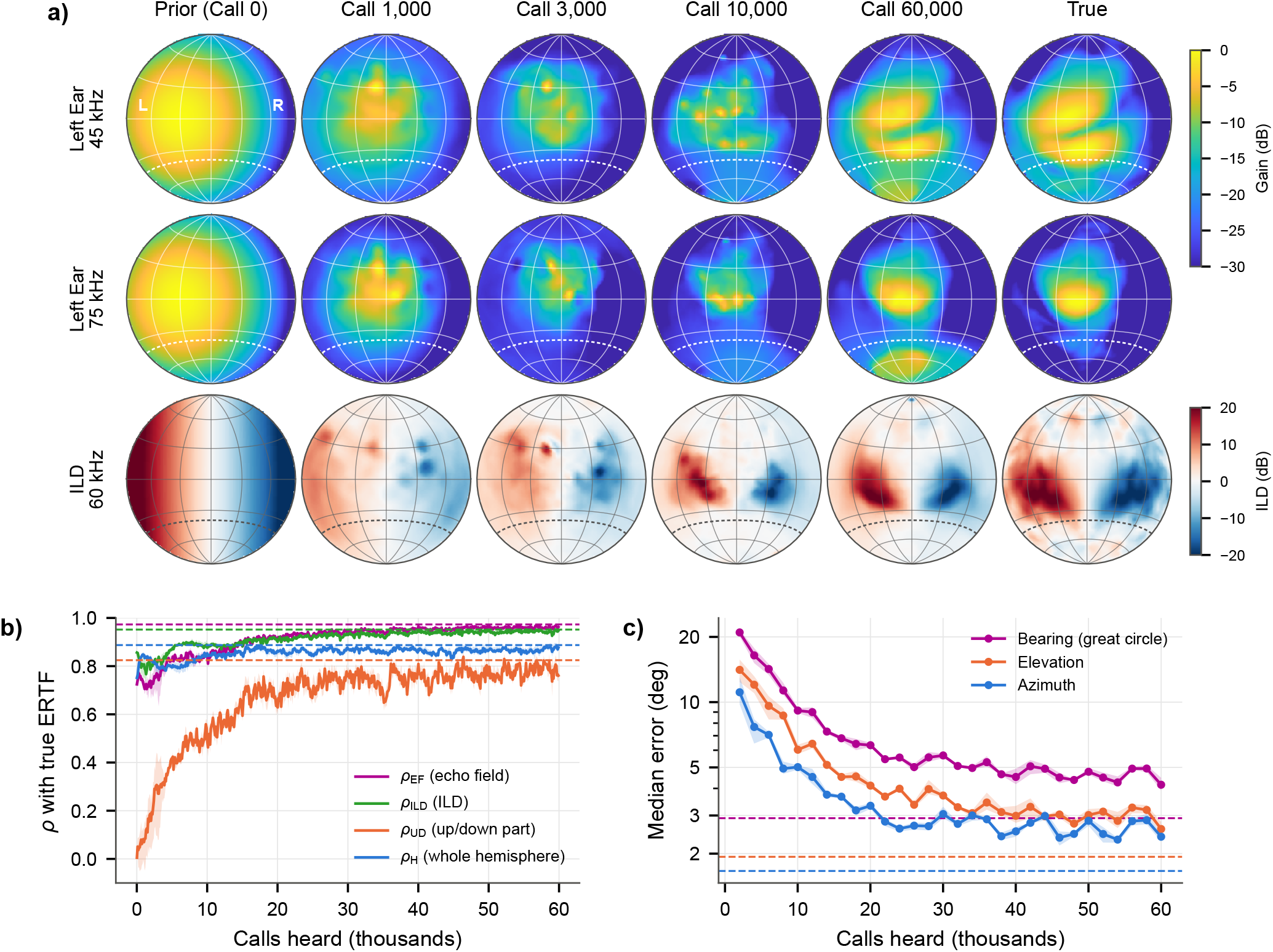
The Ears Are Learned Online, While Driving. **a**, The learned ERTF at fixed moments of the online drive, from the prior (left) to the end of the first drive (60,000 calls), next to the true ERTF (right): left ear at 45 and 75 kHz (gain in dB relative to its maximum) and the interaural level difference at 60 kHz (red: louder in the left ear). Maps of the frontal hemisphere seen from behind the head, grid every 30°. Dashed: −40° elevation, below which echoes rarely arrive. **b**, Correlation between learned and true ERTF over the drive: in the echo field (*ρ*_EF_, magenta), ILD (*ρ*_ILD_, green), up/down part (*ρ*_UD_, orange) and the whole frontal hemisphere (*ρ*_H_, blue). Dashed horizontal lines: batch-trained reference with access to the whole first drive. **c**, Median error of the direction of isolated echoes in the unseen validation worlds, inferred from a single call, over the drive (great circle, elevation and azimuth). Dashed: batch-trained reference. Lines in b and c: mean over three network seeds on the same drives, bands: range.

We score the learned ERTF against the true one only for evaluation (Fig. 2b), as the correlation *ρ* of the two directional patterns after centering them per ear and band: over the agent’s echo field, the directions from which it receives audible echoes (*ρ*_EF_), over the whole frontal hemisphere (*ρ*_H_), for the ILD (*ρ*_ILD_), and for the up/down part (*ρ*_UD_), the difference between the ERTF at mirror-image directions above and below the horizon, which carries elevation. The echo field is defined from the simulation as the directions at or above −40° elevation, from which 99.8% of the echoes arrive. This is the region where the true ERTF of both ears is within 25 dB of its maximum. It covers 35% of the frontal hemisphere (Supplementary Fig. 3). There, *ρ*_EF_ reaches 0.97 after 60,000 calls (0.964–0.969 over three seeds) and stays at 0.97 during a second drive of 60,000 calls through the same worlds (Supplementary Table 1 lists the scores of all runs). The ILD, which carries azimuth information and is largely given by the prior (*ρ*_ILD_ = 0.86), dips during the first thousand calls but recovers and ends at *ρ*_ILD_ = 0.95. The up/down part is absent from the prior. *ρ*_UD_ passes 0.5 after 6,600–7,600 calls and reaches 0.75–0.81 after 60,000 calls and 0.80 after 120,000. Over the whole frontal hemisphere, the correlation is lower (*ρ*_H_ = 0.86–0.89). Most of this gap comes from the directions below − 40°, from which only 0.2% of the echoes arrive: over all frontal directions at or above −40°, the correlation is 0.94–0.95. The directions below are hardly corrected and can even acquire spurious gain (Fig. 2a, below the dashed line).

A batch-trained reference that sees the whole first drive many times in shuffled order, and needs far more memory, reaches *ρ*_EF_ = 0.97, *ρ*_H_ = 0.89, *ρ*_ILD_ = 0.95 and *ρ*_UD_ = 0.82. The online learner comes close to it with a memory more than 700 times smaller, although the batch reference localizes isolated echoes better (2.9° against 4.4° for the main run, Fig. 2c). The weak prior speeds up the start but is not required: from a random ERTF, the agent reaches the same *ρ*_EF_ after 60,000 calls (0.960–0.968 over three seeds), only more slowly at first (0.20–0.52 instead of 0.72 after 1,000 calls, Supplementary Fig. 4).

With the learned ERTF, the agent can localize echoes in three dimensions. On the four unseen validation worlds, it points to isolated echoes (echoes without a comparable echo within *±*9 cm in range), read out at the range of each echo, from a single call with a median error of 3.9–4.4° over the seeds (2.3–2.5° in azimuth and 2.4–2.8° in elevation). It gets the sign of the elevation right for 88–94% of them, against 61° and 50% for random guesses and 59% for always answering ‘above’ (Fig. 2c, Supplementary Fig. 5). For comparison, big brown bats discriminate the elevation of targets near the horizon to within 3–4°, albeit in a different species and task [14]. During a second drive through the same worlds, the ERTF keeps improving, but the single-call bearing error drifts from about 4.5 to 5.2° (6.0° at 120,000 calls), while the ground-truth-free prediction error decreases only slightly (Supplementary Fig. 6). We therefore report localization after the first drive of 60,000 calls.

## 4 The Scene Is Learned in Three Dimensions

The EnergyScape that the agent infers from each call is a three-dimensional description of its surroundings, produced by the encoding step and trained only through the echoic prediction task. For evaluation, we placed the EnergyScapes of a whole route through an unseen world into one frame, using the simulator’s true poses, and compared their peaks with the true reflectors (Fig. 3). Of the resulting peaks, 90% are real reflectors, and they recover 32% of the reflectors heard at least five times. The matched reflectors are found with a median error of 8 cm in three dimensions and of 5 cm in height, over heights between 0 and 3 m. Information about the three-dimensional layout of the scene thus emerges along with the ERTF, as a by-product of predicting future echoes.

**Figure 3:**
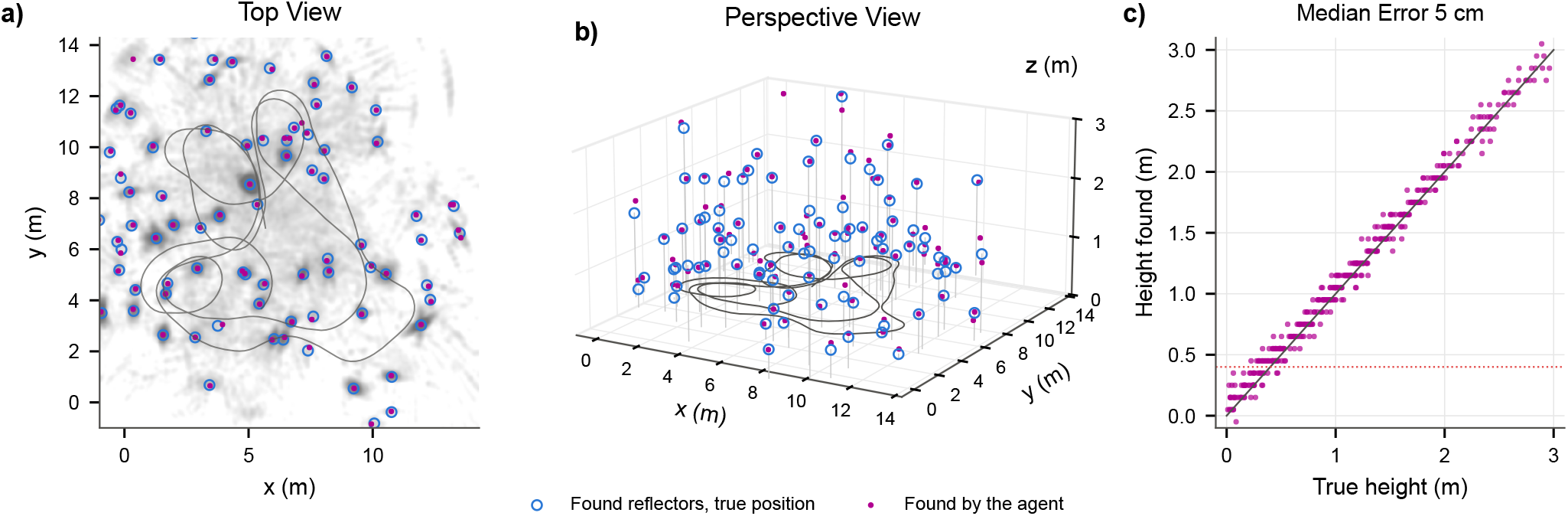
The Three-Dimensional Scene Recovered by the Agent. **a**, Top view of one unseen validation world: the reflectors the agent found, at their true positions (blue circles, reference only), and the peaks of its map (magenta), obtained by placing the EnergyScapes it inferred from single calls along its route in one frame, using the true poses for evaluation (gray: accumulated energy, gray line: route). Reflectors that were heard but not found are not shown. **b**, The same world in perspective, with the route at the height of the sensor and a stem from each found reflector to the floor (height stretched 2.6 times). **c**, Height of the found reflectors against their true height, for all matched reflectors in the four validation worlds. Red dotted line: height of the sensor. The agent’s body never leaves the horizontal plane.

## 5 Learning Requires Self-Motion

Three controls show that the learning comes from the lawful relation between the echoes and the agent’s own motion. In the first control (Fig. 4a–c), the motion readings of every training window are replaced by those of another, random moment of the drive. This replacement is done so that the motion statistics are kept but they no longer belong to the observed echoes (rather than taking random motion commands). The learning then fails, and the left/right structure of the prior is actively unlearned: *ρ*_ILD_ drops from 0.86 to −0.08, *ρ*_EF_ from 0.73 to 0.22, and the agent gets only 50% of its left/right decisions right (median error 28°). In the second control, the prediction term is removed, so that the agent only reconstructs the call it has just heard. *ρ*_EF_ then falls from 0.61 after 1,000 calls to 0.02 at the end of the drive, and the median bearing error is 47°, worse than always pointing straight ahead. In the third control, every window receives the true self-motion of the window two calls earlier, so that the motion is right in kind but misaligned by about 20 cm of travel. The ERTF is then still learned to a large part (*ρ*_EF_ = 0.87, *ρ*_UD_ = 0.64), but the agent localizes echoes poorly: the median bearing error is 19° and the up/down sign is right for only 57% of the echoes, no better than always answering ‘above’ (59%). The scene the agent infers from a call has to be moved with the motion that belongs to it.

**Figure 4:**
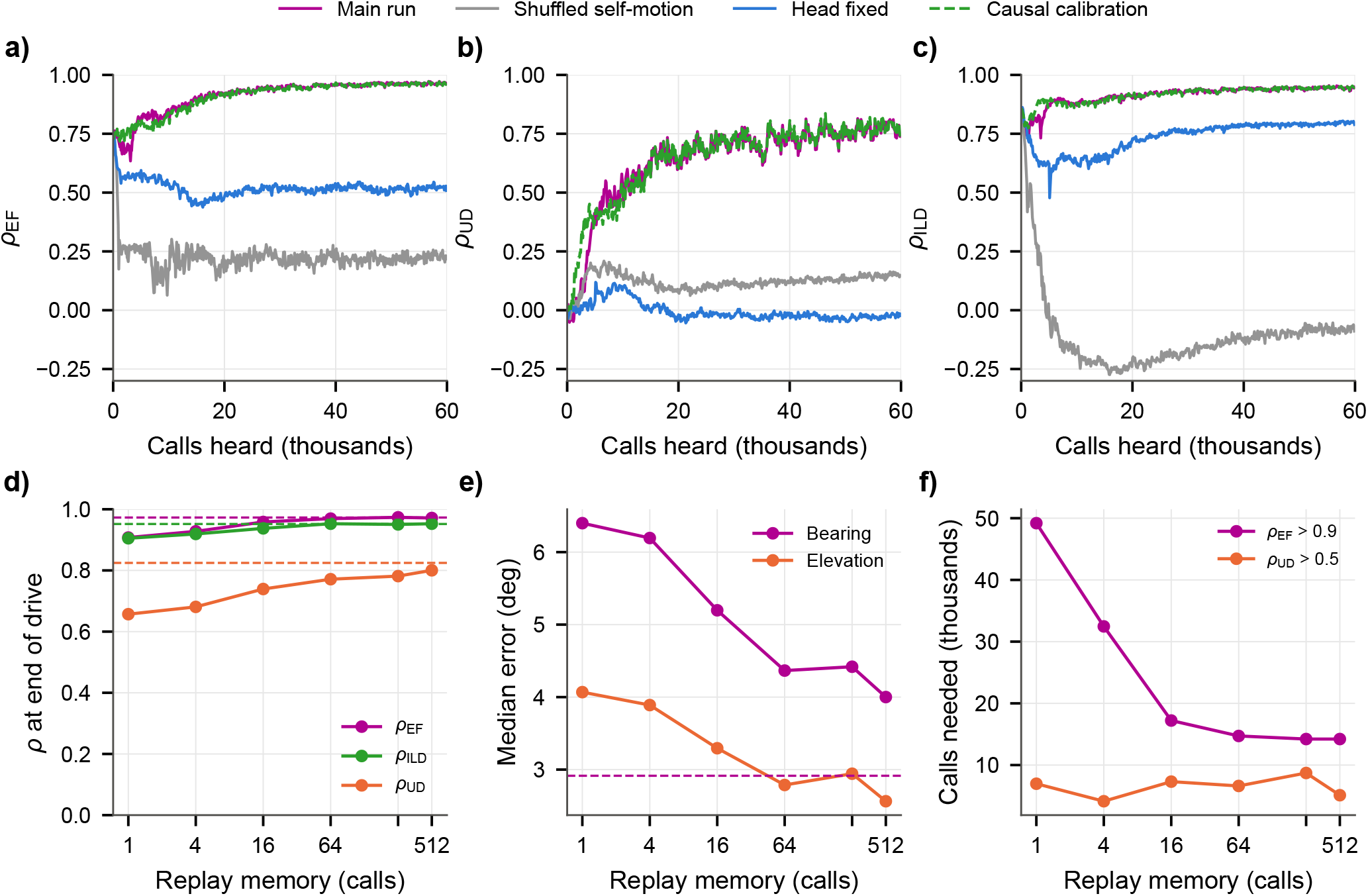
Effect of the Self-Motion and of the Replay Memory on Learning. **a**–**c**, Correlation between learned and true ERTF over the drive, in the echo field (*ρ*_EF_, a), for the up/down part (*ρ*_UD_, b) and the ILD (*ρ*_ILD_, c), for the main run (magenta), with the self-motion readings of every window taken from a random other moment of the drive (gray), with the head kept fixed (blue) and with the level scale and gain calibrated on the first 1,500 calls only (green, dashed). **d**, Correlations at the end of the drive (final model) as a function of the replay memory (calls): in the echo field (magenta), ILD (green) and up/down part (orange). **e**, Median direction error of isolated echoes in the unseen validation worlds. **f**, Calls needed to reach *ρ*_EF_ = 0.9 and *ρ*_UD_ = 0.5. Dashed in d and e: batch learner.

The learner assumes that every echo comes from a fixed point in the world. Smooth specular surfaces break this assumption: they reflect a call like a mirror [47], so that their echo comes from the foot of the surface normal, which moves along with the agent, breaking the echoic flow model. We let the agent drive twice through sparser point worlds (a sixth of the reflector density), once without surfaces and once inside a room with a floor, a ceiling and walls, with exactly the same reflectors (Supplementary Fig. 7). The surfaces carry 37% of the echo energy between 30 and 45 kHz, where the small reflectors scatter weakly, but only 9% between 75 and 90 kHz. The learned ERTF is distorted in the bands that the energy of the surfaces echoes dominate: between 30 and 60 kHz, *ρ*_EF_ falls from 0.96 to 0.86–0.88, whereas above 60 kHz it does not drop (0.85– 0.90 against 0.82–0.88), and the median bearing error grows from 6.7 to 8.2°.

## 6 Head Tilt Disentangles Elevation

When the head is kept still with respect to the body, so that the only rotations are the turns of the body about the vertical axis, the up/down part of the ERTF is not learned (*ρ*_UD_ = − 0.02), the median elevation error is 21° and azimuth is learned only partly (5.6° instead of 2.5°, Fig. 4a–c). When the head pans but neither tilts nor rolls, all motion still stays in the horizontal plane. Azimuth is then learned as well as with the full head (2.4° instead of 2.5°, left/right sign correct for 98% of the echoes), but the up/down part of the ERTF is not (*ρ*_UD_ = 0.03). Adding tilt to the head motion restores elevation: *ρ*_EF_ = 0.96, *ρ*_UD_ = 0.73 and a median bearing error of 4.1° (2.5° in elevation), with the up/down sign right for 91% of the echoes, as with the full head (0.97, 0.77, 4.4° and 88%). Motion within the horizontal plane thus leaves the sign of elevation open, and the agent settles on an arbitrary one. Without tilt, 82% of the isolated echoes come from above the sensor, but the agent places 76% of them below with the head fixed and 75% with the head panning, so that its up/down decisions are worse than always answering ‘above’ (35% and 44% correct). Rolling the head instead of tilting it does not resolve the ambiguity either. With pan and roll (roll within *±*15°), the up/down part of the ERTF is not learned (*ρ*_UD_ = 0.06), and the agent places 83% of the echoes above the sensor, so that its up/down decisions (73% correct) are no better than always answering ‘above’ (79%). The roll of our head unit is, however, smaller than its tilt (*±*15° against − 20° to +40°, a median change of 1.5° against 4.3° per call), so this comparison does not separate the axis of rotation from its amplitude. In our model, the ambiguity is resolved by tilt, in line with the role of head movements in bats [42].

The sign of elevation does not come from the layout of the scene. In our worlds, most isolated echoes (59%) come from above the sensor. When we spread the reflectors symmetrically about the height of the sensor instead (with 1.7 times as many reflectors), most come from below in the frame of the head (68%, because the head tilts up more than down). Yet the up/down part of the ERTF is learned even better (*ρ*_UD_ = 0.95 instead of 0.77, up/down sign right for 94% of the echoes, median bearing error 4.3°), as echoes now also arrive from below − 40° (5% instead of 0.2%). The sign of elevation is learned from the motion, whether most echoes come from above or from below.

## 7 A Short Memory Suffices

The online learner keeps only a replay memory of *N* calls, a training windows that end within its last *N* + 1 calls (Methods). Fig. 4d–f shows how the result depends on *N*. Going from 64 to 512 calls, eight times the memory, raises *ρ*_EF_ only from 0.969 to 0.971 and *ρ*_UD_ from 0.77 to 0.80, and the calls needed to reach *ρ*_EF_ = 0.9 stay at about 14,000. With *N* = 16, *ρ*_EF_ reaches 0.96 (0.9 after about 17,000 calls). Even with *N* = 1, where every step uses only the newest windows, it reaches 0.91 and the median bearing error is 6.4° instead of 4.4°, although *ρ*_EF_ = 0.9 is only reached after about 49,000 calls. A memory of 64 calls comes close to the batch learner that stores the whole drive of 60,000 calls.

## 8 Changed Ears Are Relearned Quickly

Ears can change during a bat’s life, for example as they grow or after damage. To mimic such a change, we rotated the directivity of the left ear upward and that of the right ear downward by 5, 10 or 20°, leaving the emitter as it was, after the agent had completed its 120,000-call drive, and let it continue to drive and learn with the same short-term memory setup (Fig. 5). For a rotation of *±* 10°, *ρ*_EF_ drops from 0.97 to 0.91 and *ρ*_UD_ to 0.68. After about 200 calls (some 20 m of travel, when the 200-step warm-up of the learning rate ends), the agent’s ERTF agrees better with its new ears than with its old ones, and after about 1,000 calls it has recovered 90% of the way. Starting from the same *ρ*_EF_, the first learning needed about 31,000 calls to gain as much. A rotation of *±*20° causes a larger drop (to *ρ*_EF_ = 0.79 and *ρ*_UD_ = 0.48), yet the agent recovers 90% of the way within about 700 calls (first learning from the same *ρ*_EF_: about 27,000 calls), and for *±*5° the drop is small (to 0.95) and is undone within about 1,000 calls as well (first learning: about 26,000 calls). Relearning is 25 to 38 times faster than the first learning. After the larger rotations, *ρ*_UD_ ends above its value before the change (0.88 at *±* 10° and 0.94 at *±* 20°, against 0.80). We do not read this as better hearing: rotating the two ears in opposite directions makes up and down differ more strongly in the ERTF itself, which makes this part easier to recover. Humans, too, adapt to altered spectral cues quickly when they receive intensive sensorimotor training, within about 20 minutes [48].

**Figure 5:**
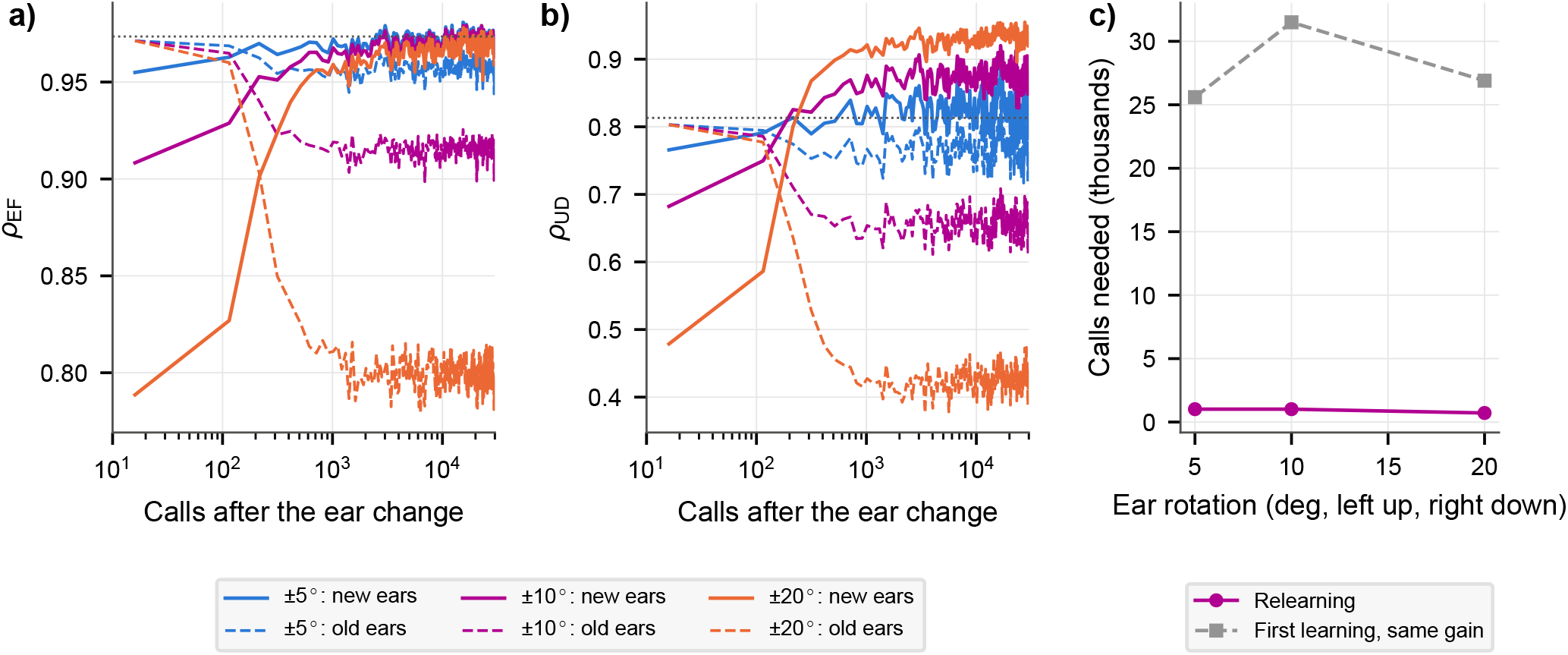
Relearning After a Change of the Ears. At call 0, the left ear’s directivity is rotated upward and the right ear’s downward by *±*5, *±*10 or *±*20°, while the emitter is unchanged. The agent keeps driving and learning with the same replay memory (*N* = 64). **a**, Correlation in the echo field (*ρ*_EF_) with the new ears (solid) and with the old ears (dashed), against the number of calls since the change (logarithmic). Dotted: *ρ*_EF_ before the change. **b**, The same for the up/down part (*ρ*_UD_). **c**, Calls needed to recover 90% of the way to *ρ*_EF_ before the change, against the size of the rotation (magenta), next to the calls the first learning needed to gain as much from the same starting value (gray).

## 9 Discussion

We have shown that an echolocating agent can learn its own ERTF in azimuth and elevation without an external teacher, using echoacoustic flow, by requiring that the scene it infers from one call predicts the echoes of the next calls. It does so online, from a memory of about 80 calls. Along the way, its scene estimates come to encode the positions of reflectors in three dimensions, and it relearns the spectrospatial mapping after the ears change. In contrast to earlier sensorimotor accounts of auditory space [9, 26], the agent learns the transfer function of its ears itself, not a mapping from cues to directions, and it exploits the fact that an echolocator is its own sound source: every call returns echoes from a static world whose apparent motion is fully determined by the agent’s own movement. This extends our earlier work on acoustic flow [29, 34] and sonar array self-calibration [36] to a biologically plausible binaural head in three dimensions.

The agent has to survive while its model of its ears is still wrong. Range follows from the delay of the first echo, and the sign of the ILD, which the weak prior already provides, is enough to steer away from obstacles without localizing them [44, 45]. Such cues would let an animal move safely while it learns its ERTF, although our agent follows prescribed paths and does not show this. Young bats may even start the calibration before they fly. Big brown bat pups fly only after about a month [49], but they already emit echolocation call sequences before they achieve powered flight [50], so that head movements in the roost could provide the first self-motion to learn from. We state this as a hypothesis.

Our results lead to predictions that can be tested in bats. First, if bats use a comparable mechanism, restraining the out-of-plane rotations of the head and body (called tilt in our model) should impair the relearning of elevation after an ear manipulation, while azimuth is largely relearned. Second, the ERTF should be learned best in the echo field, so that animals raised in environments with different vertical structure should localize best in different directions. Third, after a manipulation such as a tragus deflection [15, 16], localization in elevation should recover gradually with the number of calls emitted in flight, rather than with time alone. Fourth, bats that calibrate among smooth surfaces, such as walls or water, should localize less well with the frequency bands in which these surfaces dominate the echoes.

The present study has several limitations. For one, all results are obtained in simulation, mostly in worlds of point reflectors. However, the simulation environments are well calibrated to real-world environments and have been shown to generalize well to real-world acoustic scenarios (Methods). The learner also assumes a static scene of point-like reflectors. Specular surfaces break this assumption and distort the ERTF in the bands they dominate, and moving prey does not follow the agent’s self-motion. However, prey echoes are typically less prominent in the echoic observations of bats and contribute relatively little to the overall echo energy. Another shortcoming is that the self-motion is known to the agent up to the specified observation noise. For this information, a bat has to rely on its vestibular system and its motor commands, which may suffer from inaccuracies in practice. Pose drift over time is not a problem, as only short windows are considered in the learning process. A final shortcoming is that the agent does not choose its movements to learn faster, which bats, with their active pinnae and head movements, may well do. We do not claim that our architecture is the one implemented in the bat’s brain. Within the assumptions above, the prediction of self-generated echoes under self-motion suffices to learn the ERTF.

## 10 Methods

### Simulated Sonar Agent and Worlds

All experiments were carried out in simulation, on a ground robot carrying a bat-like sonar head (one emitter, two ears) on a pan-tilt-roll unit. Except in experiment E9 (experiments are denoted as Ex, see below), the worlds fulfill the learner’s central assumption, that every echo comes from a fixed point in the world: they consist of isotropic point reflectors only, without walls, floors or other specular surfaces. The side of a square arena is drawn uniformly between 12 and 16 m, and reflectors are scattered uniformly (Poisson count, 0.3 per m^3^) in a box that extends 3 m beyond the arena on every side and from the floor up to 3 m, giving 284 to 448 reflectors per world, with random backscatter strengths *σ*_*n*_, sizes and spectral tilts (Supplementary Note 3). We generated 40 training and 4 validation worlds from different random seeds. Acoustic simulations of this kind have been calibrated against, and shown to transfer to, real-world measurements for biologically relevant ensonifications [51, 52]. The symmetric worlds of E8 spread the reflectors over 0.40 *±* 2.6 m in height, about the height of the sensor. The worlds of E9 have a sixth of the reflector density (0.05 per m^3^), without surfaces or inside a room of specular surfaces with the same reflectors: a floor, a ceiling at 3.2 m and four walls 3 m beyond the arena, whose echoes come from the foot of the perpendicular from the sensor.

Directions *ψ* = (*θ, φ*) consist of the azimuth *θ* (positive to the left) and the elevation *φ* (positive upward) in the frame of the sonar head, with unit vector 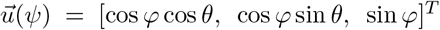. The directivities of the emitter, *H*_*E*_(*ψ, ω*), and of the ears, 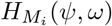 with *i* ∈ {1, 2}, are those of the simulated head-related transfer function of *Phyllostomus discolor* [3] (amplitude gains on a 2.5° grid, 141 frequencies between 25 and 95 kHz, interpolated bilinearly in direction and linearly in frequency). This transfer function is used only to synthesize the echoes and to score the result. It never enters the learning. The call *s*_*b*_[*k*] is a 3 ms linear frequency-modulated downsweep from 90 to 30 kHz, shaped in magnitude to the call spectrum supplied with the HRTF data set, with 5% raised-cosine ramps. It is a simplification of the multi-harmonic calls of *P. discolor*, and only the ERTF is species-specific. For the head pose of a call, we synthesize the signal received by ear *i* coherently in the frequency domain:

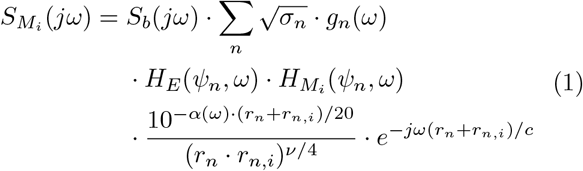

where *S*_*b*_(*jω*) is the DFT of the call, *r*_*n*_ the distance from the emitter to reflector *n, r*_*n,i*_ the distance from the reflector to ear *i* (the ears sit at *±*1.2 cm from the emitter on the interaural axis, so interaural time differences are included), *ψ*_*n*_ the reflector’s direction, *g*_*n*_(*ω*) its roll-off and tilt, *α*(*ω*) the atmospheric absorption in dB/m (ISO 9613-1, 20 °C, 50% relative humidity [53]), *c* = 343 m/s, and *ν* the spreading exponent of the echo power. All experiments reported here used *ν* = 2, a weaker spreading than the physical *ν* = 4 of a point reflector, for both the synthesis and the learner. The sum runs over the reflectors in front of the head between 1.0 and 10.5 m. Reflectors closer than 1 m are dropped, which emulates a receiver that is blanked during and shortly after the call. This synthesis is deliberately richer than the learner’s model, which ignores echo interference, interaural time differences and reflector-specific spectra. The world is static and the head does not move during a call.

### Head and Body Motion

The body drives in the plane of the floor at a constant 0.10 m per call, without turns on the spot or reversing. Its path curvature follows an Ornstein–Uhlenbeck process (correlation length 1.5 m, stationary standard deviation about 0.5 m^*−*1^, limited to a turning radius of at least 0.8 m), with a term that steers the body back toward the arena’s center within 2.5 m of its edge. The head unit has pan *χ*_*p*_, tilt *χ*_*t*_ and roll *χ*_*r*_ (positive to the left, up, and left ear up), applied in that order, which places the head in the body frame as follows:

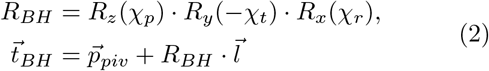

where *R*_*x*_, *R*_*y*_ and *R*_*z*_ are rotations about the forward, left and upward axes, 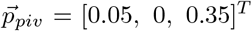 m is the pivot of the head unit in the body frame and 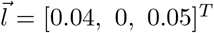 m the lever arm from the pivot to the sensor (0.40 m above the floor at zero tilt). The joints are limited to *±* 60° (pan), − 20° to +40° (tilt) and *±*15° (roll). The joint angles follow smooth random scanning signals (Supplementary Note 3), with a median absolute change per call of 2.9° (pan), 4.3° (tilt) and 1.5° (roll). The learner receives only what the robot itself measures: the body displacement between calls from wheel odometry (10% multiplicative plus 1 mm additive Gaussian noise per component), the yaw increment from a gyroscope (0.15° noise) and the three joint angles from the servos (0.2° noise each). The true poses are used only for evaluation. Each world is traversed with 1,500 calls (150 m) for training and 1,000 calls for validation, so that one drive through the 40 training worlds, in a fixed order, contains 60,000 calls (6 km). Further drives through the same worlds, with new routes, head motion and noise, were generated by offsetting all random seeds.

### Cochlear Model

Each received signal is converted into a cochleogram [12, 54], using *C* = 64 bands with center frequencies spaced logarithmically between 30 and 90 kHz, each a Gaussian *H*_*c*_(*ω*) on a logarithmic frequency axis with a full width at half maximum of two band spacings. Combined with a matched filter, the cochleogram of ear *i* reads:

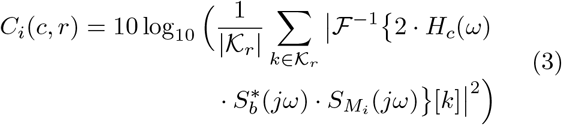

where the one-sided inverse DFT ^*−*1^ yields the analytic band signal, 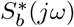 is the matched filter (normalized to unit peak gain), and _*r*_ holds the samples whose two-way travel time 2*r/c* falls into range cell *r*. We use 242 cells of 3 cm between 0.25 and 7.51 m, so one call yields a [2 *×*64 *×*242] cochleogram. It is converted into normalized levels:

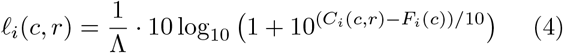

with *F*_*i*_(*c*) the noise floor of ear *i* and band *c* (Supplementary Note 3) and Λ = 41.0 dB the 99.9th percentile of the numerator. Both come from the cochleograms alone, computed over the whole first drive (over the first 1,500 calls only in C5).

### EnergyScape

From a single call *t*, an encoder infers the EnergyScape, a description of the scene around the head. It is defined on a grid of range cells *r* and directions *ψ*_*j*_ in the head frame. The directions are the *D* = 1,407 vertices of a level-4 icosphere (about 4°apart) within 96° of straight ahead, which cover the frontal hemisphere with a small margin, each with a quadrature weight *w*_*j*_, its share of the surrounding spherical triangles. Every cell of the grid holds two quantities. The first is the reflected energy *P*_*t*_(*r, ψ*_*j*_). The second is a set of four color weights *π*_*m*_(*r, ψ*_*j*_), which sum to one and describe the spectrum of the echo from that cell as a mixture of four learned echo spectra *B*_*m*_(*c*). These spectra are shared by all cells and all calls.

The encoder treats the cochleogram as a signal over range with 2 *· C* = 128 channels: a one-dimensional convolutional network gives, per range cell, the real spherical-harmonic coefficients of the energy (up to degree 32) and of four color logits (up to degree 12). On the icosphere, a softplus turns the energy into 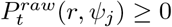, starting close to zero so that the agent initially hears nothing, and a softmax over the logits gives the color weights *π*_*m*_(*r, ψ*_*j*_). A learned blur on the sphere and a Gaussian of one cell in range then give *P*_*t*_(*r, ψ*_*j*_). Supplementary Note 1 describes the layers.

### Acoustic-Flow Warp

The EnergyScape inferred from call *t* has to predict the cochleograms of the calls that follow. As the world is static, the self-motion fixes where each part of the scene appears at a later call. The cell (*r*_*i*_, *ψ*_*j*_) in the head frame of call *t* + *k* holds the same world point as the point 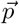 in the head frame of call *t*:

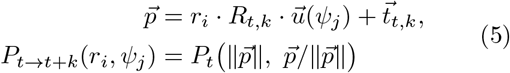

where 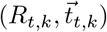 is the pose of the head at call *t* + *k* in the head frame of call *t*. This pose is obtained by integrating the odometry and gyroscope increments and composing them with the measured joint angles through equation (2). The color weights *π*_*m*_ are moved in the same way. *P*_*t*_ and *π*_*m*_ are sampled linearly in range and barycentrically on the icosphere. Points outside the range grid or the field of view contribute nothing. The warp has no learned parameters and is exact for any step size. Its small-step limit is the continuous acoustic flow [29, 34].

### Differentiable Renderer

A renderer with prescribed spreading and a learned sensor model turns the moved EnergyScape into the levels expected at call *t*+*k*, for ear *i*, band *c* and range cell *r*:

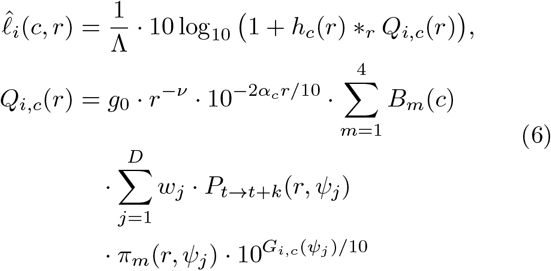

where *π*_*m*_ are the moved color weights, *G*_*i,c*_(*ψ*) is the learned ERTF in dB, *α*_*c*_ a learned one-way absorption (initialized at 0.035 dB/m per kHz), *h*_*c*_(*r*) a learned 13-tap range point-spread function per band, *\**_*r*_ a convolution over range, *ν* the spreading exponent of the data, and *g*_0_ a fixed gain. We calibrate *g*_0_ from the cochleograms alone (Supplementary Note 3). Fixing *g*_0_ removes the trade-off between the level of *P* and the gain of the ERTF. Every step of the warp and the renderer is differentiable, so the prediction error is passed back to the ERTF and, through the warp, to the encoder.

### ERTF Model and Priors

As emitter and ear directivities always appear as a product in an echo (equation (1)), what can be learned is their product, the ERTF per cochlear band. A small multilayer perceptron *f* represents it for both ears and all bands:

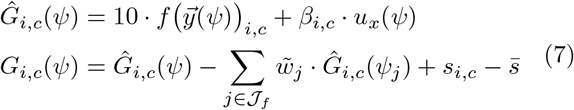

where 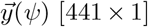 holds the real spherical harmonics up to degree 20 (pole on the forward axis), *f* has two hidden layers of 128 units (GELU) and 2 · *C* = 128 outputs, *β*_*i,c*_ is a learned forward-beam term (initialized at zero) acting through the forward component *u*_*x*_(*ψ*) of 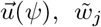 are the quadrature weights normalized over the frontal hemisphere *J*_*f*_, and *s*_*i,c*_ is a direction-independent system term with mean 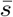 over ears and bands. The directional part thus has zero mean over the frontal hemisphere, and the overall level is set by *g*_0_. Nothing in the model encodes symmetry, handedness or an up/down cue. Which parts of the ERTF the prediction can pin down depends on the motion: a mirrored scene with a mirrored ERTF predicts the same echoes for any motion that maps onto itself under the mirror. The head-axis experiments (E7) test this. Apart from the prior below, the learner is given the call (as the matched filter), the spreading exponent *ν* of the data, the form of the absorption, the assumption that the world is static, and the measured self-motion. It is not given the emitter directivity, the ear directivities or the direction of any echo. Before learning, *f* is fitted (3,000 Adam steps) to a weak prior that states only that one ear points left and the other right:

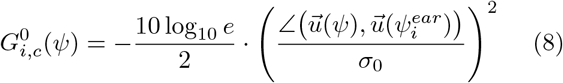

with 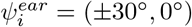 (left ear positive), *σ*_0_ = 30° and the same lobe in every band. This prior is symmetric in elevation, so its up/down part is zero. It is not enforced afterward. Fitted to the true ERTF instead, as a check, the same network reaches a correlation of 0.9998 in the echo field (Supplementary Fig. 8). The complete model has 1,479,593 parameters (Supplementary Fig. 1).

### Training Objective

The model is trained on windows of 17 consecutive calls *t*, …, *t* + 16 within one world, together with the motion readings between them. The encoder sees only call *t*. Per window, the loss reads:

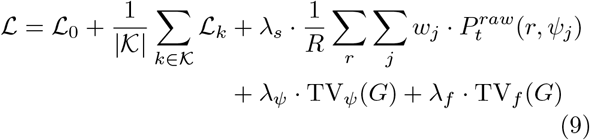

with the following terms:

- *Prediction error ℒ*_*k*_: the mean squared error between the levels predicted for call *t* + *k* (equations (5) and (6)) and the levels heard at that call, over both ears, all bands and the range cells beyond 1.06 m (receiver blanking). It is averaged over *k* ∈ *K* {= 1, 2, 3, 4, 6, 8, 12, 16}. This term carries the self-motion into the learning.
- *Reconstruction error ℒ*_0_: the same error for call *t* itself, which the warp leaves in place (*k* = 0).
- *Sparsity*: an ℒ_1_ penalty on the EnergyScape before blurring, 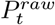, weighted by solid angle and averaged over the *R* range cells (*λ*_*s*_ = 1). As 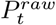 is positive (softplus), no absolute value is needed. The penalty keeps the inferred scene compact.
- *Smoothness of the ERTF* : two smoothed total-variation norms with *ϵ* = 1 dB, which penalize steep edges only linearly, so that spectral notches can survive [55]:
  – across directions, TV_*ψ*_(*G*): the solid-angle-weighted mean of 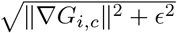 (in dB/rad), with the gradient taken of the piecewise-linear interpolant on the icosphere’s triangles, averaged over ears and bands (*λ*_*ψ*_ = 10^*−*5^).
  – across bands, TV_*f*_ (*G*): the mean of 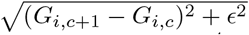 over neighboring bands (*λ*_*f*_ = 3 *·* 10^*−*4^).

Ground truth enters none of these terms, so the learning is self-supervised. We use AdamW [56] with learning rate 3 *·* 10^*−*4^, weight decay 0.01, gradient norm clipped at 1, 16 windows per step and a linear warm-up over 1,000 steps.

### Online Learning

In the online experiments, the agent learns while it drives through the worlds. The training worlds are traversed one after the other, and for every new call *p*, one optimization step is taken on a minibatch of 16 windows drawn uniformly, with replacement, from a replay memory of *N* calls, that is, from the windows whose last call lies among the most recent *N* + 1 calls:

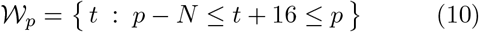

where *t* is the first call of a window. A memory of *N* calls thus holds the *N* + 1 newest windows, with the data of the last *N* + 17 calls. After the warm-up, the learning rate stays constant, as an online learner has no end of training to decay toward. The main online run (E1) used *N* = 64 (81 calls of raw history, 8 m of travel). After the first drive (60,000 calls, 59,984 steps), it continued for a second 60,000-call drive through the same worlds with new routes, head motion and noise, starting from the final weights (fresh optimizer state, 200-step warm-up, level scale and gain of the first drive), for 120,000 calls in total. As a reference (E2), we trained the same model in batch mode on all 59,360 windows of the first drive, in shuffled epochs under a cosine schedule, until its ground-truth-free validation prediction error stopped improving clearly (64,000 steps, Supplementary Note 3). We report its checkpoint with the lowest validation prediction error. E2 saw only the first drive.

### Further Experiments

All start from the setup of E1 (Supplementary Note 2 lists all experiments):

- *Relearning (E3)*: we rotated the directivity of each ear about the interaural axis, the left ear up and the right ear down by *δ ∈ {* 5°, 10°, 20°*}*, leaving the emitter unchanged. Online learning (*N* = 64) continued for 30,000 calls on a new drive from the end of the 120,000-call drive of E1 (fresh optimizer state, 200-step warm-up, E1’s level scale and gain). The calls needed to relearn are those until *ρ*_EF_ has recovered 90% of the way from its value just after the change to its value at the end of E1.
- *Replay memory (E4)*: we repeated the first drive of E1 with *N ∈ {* 1, 4, 16, 256, 512*}* calls (E1 provides *N* = 64).
- *Head axes (E7)*: the first drive of E1 with pan only, with pan and tilt, or with pan and roll, on the same body routes.
- *Symmetric worlds (E8)* and *specular surfaces (E9)*: one drive of 60,000 calls through the worlds described above.

### Controls and Ablations

Each repeats the first drive of E1 with one change (Supplementary Note 2):

- *Shuffled motion (C1)*: the self-motion readings of every window come from a random other window of the drive.
- *No prediction (C2)*: only the reconstruction of the current call is learned.
- *Shifted motion (C3)*: every window gets the true self-motion of the window two calls earlier.
- *Head fixed (C4)*: no pan, tilt or roll, on the same body routes.
- *Causal calibration (C5)*: level scale and renderer gain from the first 1,500 calls only.
- *No prior (E6)*: *f* starts from its random initialization instead of the prior.
- *No smoothness (E6)*: *λ*_*ψ*_ = *λ*_*f*_ = 0.

### Evaluation

Ground truth was used only for evaluation. The true ERTF is the product of the emitter and ear power gains, averaged over each band with its power response (including the matched filter), on the 1,313 icosphere directions of the frontal hemisphere. True and learned ERTF are centered per ear and band, and all statistics are weighted by solid angle. End-of-drive scores are those of the final model (*ρ*_UD_ fluctuates by about *±*0.04 between snapshots 100 calls apart). We report the Pearson correlation *ρ* over all frontal directions, ears and bands (*ρ*_H_), and over the echo field (*ρ*_EF_), the directions from which echoes arrive: within 25 dB of the maximum gain for both ears and at elevations of at least −40° (455 directions, 35% of the frontal solid angle, with both patterns re-centered on this region). Of the 7.3 *·* 10^6^ echo sources listed by the simulator during the first drive, 99.84% arrived from − 40° or higher. Within 25 dB, we also report the correlation of the interaural level difference *G*_1,*c*_ −*G*_2,*c*_ (*ρ*_ILD_), of the up/down part (*ρ*_UD_) *G*_*i,c*_(*θ, φ*) − *G*_*i,c*_(*θ, − φ*) for *φ >* 3°, which the prior lacks and which pairs every direction above +40° with a rarely heard one below − 40° (restricted to pairs within *±*40°, *ρ*_UD_ = 0.91–0.93 for E1), and of the left/right part *G*_*i,c*_(*θ, φ*) − *G*_*i,c*_(*θ, φ*).

Localization was scored on isolated echoes in the validation worlds (every third call, every sixth for curves over the drive, criteria in Supplementary Note 3). The agent’s bearing is the frontal direction that maximizes *P*_*t*_(*r, ψ*), weighted by the learned mean gain and summed over *±*2 cells around the echo’s range. We report median great-circle, azimuth and elevation errors and the fraction of correct up/down and left/right decisions (|*φ*|, |*θ*| *>* 5°), next to the errors of uniformly random frontal directions and the share of correct up/down decisions obtained by always answering ‘above’.

For the scene, the gain-weighted EnergyScapes of a validation route were placed in the world frame with the true poses, for scoring only, and their local maxima were matched to the reflectors heard at least five times (Supplementary Note 3). We report recall, precision and the median three-dimensional and height errors over the four validation worlds.

### Statistics and Reproducibility

The first drive of E1 was run with three random seeds (0, 1 and 2, E5), which change the network initialization, the prior fit and the replay sampling but not the simulated drives, and so was the run without the prior (E6). We report the mean and the range over seeds. The other experiments were run once (seed 0). Supplementary Note 2 lists all experiments. Localization errors are medians over 3,990 isolated echoes in the four unseen validation worlds. No score was ever fed back into training, model selection used only the ground-truth-free validation prediction error, and no hypothesis tests were performed. The software was written in Python 3.13 with PyTorch 2.11 and run on NVIDIA L40S and RTX PRO 5000 GPUs. One 60,000-call online drive takes about 3.7 h on one GPU.

### Use of AI Tools

The AI assistant Claude (Anthropic) was used to implement the simulation, learning and plotting software, to run and monitor the experiments under the author’s instructions and to draft parts of the text. The author checked all results, reviewed and edited the text and takes full responsibility for the content of this article.

## Data Availability

The simulated worlds, and for every experiment the configuration, the final model, the learned ERTF over the drive and the evaluation metrics, as well as the source data of the figures, are available at Zenodo (https://doi.org/10.5281/zenodo.23537553). Supplementary Video 1 is available at https://youtu.be/6nxHsedi9fE and in the Zenodo record. The cochleograms, motion readings and evaluation ground truth are regenerated exactly from the seeds with the code. The *Phyllostomus discolor* HRTF is that of ref. [3] and is included in the Zenodo record with the permission of its authors.

## Code Availability

The simulator, training, evaluation and figure code and the exact run configurations are available at Zenodo (https://doi.org/10.5281/zenodo.23537553) under the MIT license.

## Author contributions

J.S. conceived the study, developed the model, carried out the experiments and wrote the paper.

## Additional information

Supplementary Information is available for this paper. Correspondence and requests for materials should be addressed to Jan Steckel. Reprints and permissions information is available at www.nature.com/reprints.

## 11 Supplementary Information

### 11.1 Supplementary Note 1: The Model Architecture

This note describes the model layer by layer (Supplementary Fig. 1). The model has three parts: an encoder that infers the scene from a single call, a warp that moves this scene with the measured self-motion, and a renderer that predicts the cochleograms of later calls through the learned ERTF. Only the encoder, the width of the spherical blur, the ERTF network and three small sets of renderer parameters are learned. The warp and the spreading and gain of the renderer are fixed. The complete model has 1,479,593 parameters, of which 1,388,582 are in the encoder, 3 in the blur, 89,856 in the ERTF network and 1,152 in the rest of the renderer.

#### Input

The input is the cochleogram of one call in normalized levels ℓ_*i*_(*c, r*) (Methods, equation (4)): two ears, 64 bands between 30 and 90 kHz and 242 range cells of 3 cm between 0.25 and 7.51 m. The encoder treats it as a signal over range with 2 *×*64 = 128 channels, so that every layer sees both ears and all bands at once and works along range only.

#### Encoder

An input layer (a one-dimensional convolution from 128 to 192 channels with kernel size 5, followed by group normalization with 8 groups and a GELU) is followed by five residual blocks. Each block applies a convolution from 192 to 192 channels with kernel size 5, group normalization and a GELU, and adds the result to its input. The dilations of the five blocks are 2, 4, 1, 2 and 4, so that the receptive field of an output cell spans 57 range cells, or about 1.7 m. A final 1 *×* 1 convolution maps the 192 channels of every range cell to 1,765 numbers: the 1,089 coefficients *a*_*l*_(*r*) of a real spherical-harmonic expansion up to degree 32 of the EnergyScape, and 4 *×* 169 coefficients *b*_*m,l*_(*r*) up to degree 12 of four color logits. Its weights are initialized with a standard deviation of 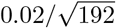 and its biases at zero, so that the initial scene is close to empty. The encoder runs in bfloat16 precision. Everything after it runs in single precision.

#### From Coefficients to a Scene on the Sphere

The coefficients are evaluated on the *D* = 1,407 frontal vertices *ψ*_*j*_ of a level-4 icosphere (about 4° apart), with *Y*_*l*_ the real spherical harmonics. The EnergyScape before blurring and the color weights are

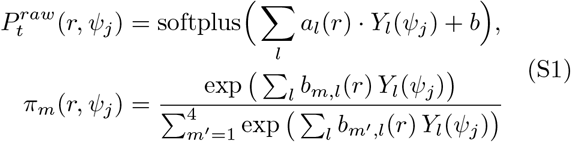

with a learned bias *b* initialized at − 15, so that the agent starts by hearing nothing. 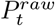 is then blurred on the sphere by a heat kernel, applied exactly in the spherical-harmonic domain. With 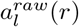 the spherical-harmonic coefficients of 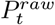 and *l* the degree of *Y*_*l*_, the blurred coefficients and the width of the kernel are

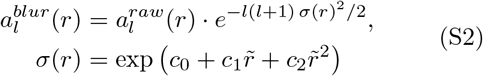

where 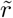 is the range scaled to [0, 1]. The width has three learned parameters, starts at 3° and is limited to 0.5– 25°. A Gaussian with a standard deviation of one cell, truncated at *±*3 cells, blurs the result in range. This gives the EnergyScape *P*_*t*_(*r, ψ*_*j*_), an array of 242 *×* 1,407 cells, and the color weights *π*_*m*_, an array of 4 *×* 242 *×* 1,407.

#### Warp

For every later call *t* + *k* with *k ∈ {*1, 2, 3, 4, 6, 8, 12, 16*}*, the pose of the head at call *t* + *k* in the head frame of call *t* is computed from the measured self-motion: the wheel odometry and the gyroscope give the body poses, and the joint angles give the head in the body (Methods, equation (2)). Each cell (*r*_*i*_, *ψ*_*j*_) of the frame of call *t* +*k* is mapped to the point 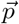 in the frame of call *t* (Methods, equation (5)), and *P*_*t*_ and *π*_*m*_ are read there, linearly between range cells and barycentrically within the spherical triangle that contains the direction 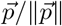. Points outside the range grid or the frontal field of view read zero. The warp has no parameters and is exact for any size of the step. The case *k* = 0 is the identity and gives the reconstruction of call *t* itself.

#### ERTF Network

The ERTF is a function of direction, the same for every call. Its input is the vector of the 441 real spherical harmonics up to degree 20 of the direction (pole on the forward axis, scaled by 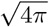). Two hidden layers of 128 units with GELU activations lead to an output layer of 2 *×* 64 = 128 values, one per ear and band, which is multiplied by 10 to give decibels. The output layer starts with weights of standard deviation 0.01 and zero biases. A learned forward term, the mean over the frontal hemisphere and a system term complete the ERTF:

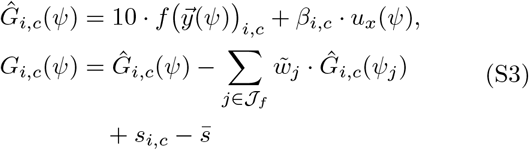

The forward term *β*_*i,c*_ *u*_*x*_(*ψ*) depends only on the angle from the forward axis (128 parameters, initialized at zero). The second term subtracts the mean over the frontal hemisphere *J*_*f*_, weighted by solid angle, per ear and band. The system term 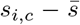 (128 parameters) does not depend on direction. This fixes the gauge: the directional part has zero mean, and the overall level is set by the fixed gain *g*_0_ of the renderer rather than traded against the level of the EnergyScape (Methods, equation (7)). Before learning, the network is fitted to the weak prior (Methods, equation (8)) with 3,000 Adam steps (learning rate 3 *·* 10^*−*3^, cosine decay), weighted by solid angle over the frontal hemisphere. Evaluated on the icosphere, the network gives the ERTF *G*_*i,c*_(*ψ*_*j*_), an array of 1,407 *×* 2 *×* 64 values in dB.

#### Renderer

For each ear *i*, band *c* and range cell *r*, the renderer sums the energy of the moved EnergyScape over directions, weighted by the quadrature weights *w*_*j*_, the color of each cell and the ERTF, and applies spreading and absorption:

**Supplementary Fig. 1:**
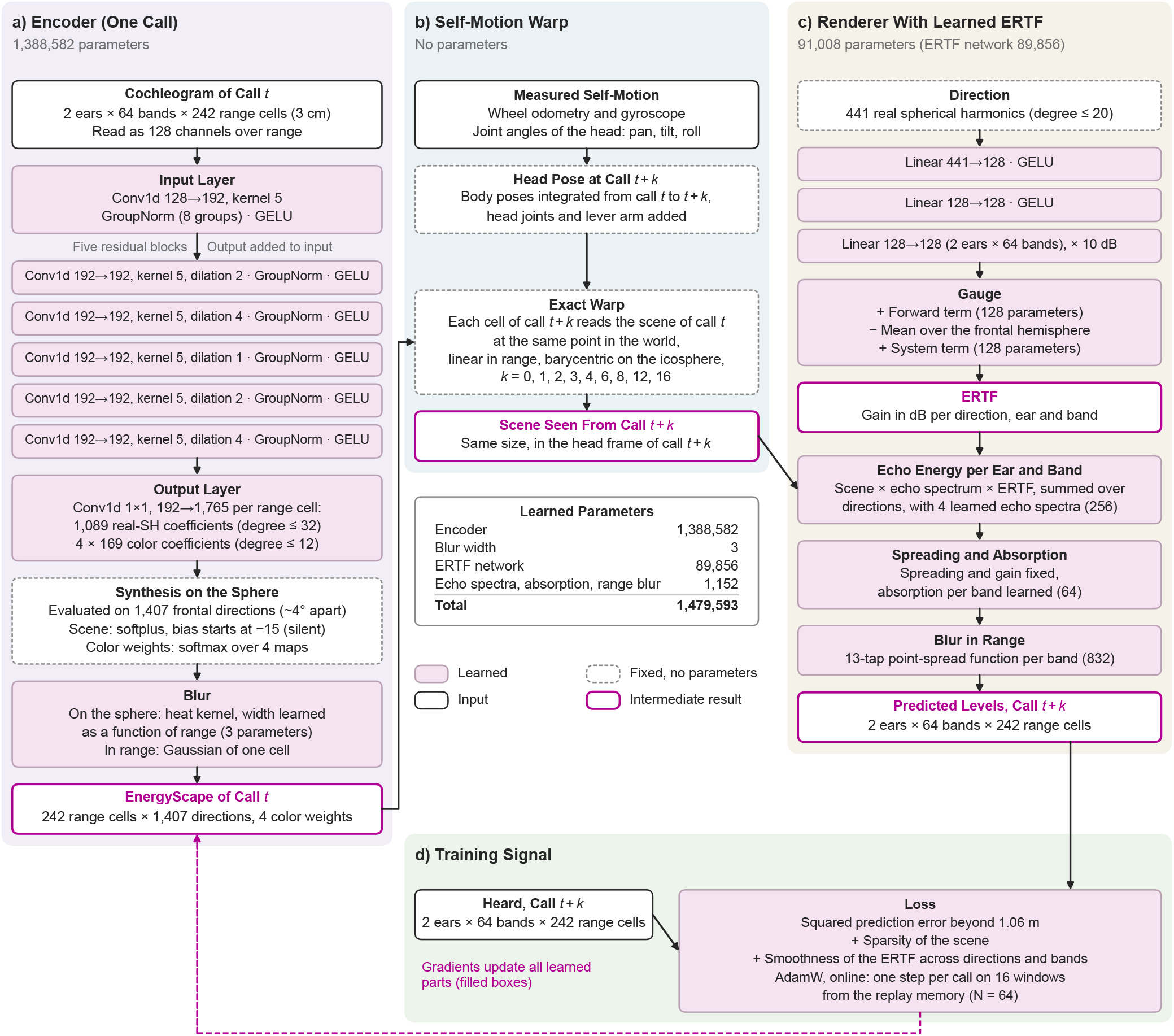
The Model in Detail. **a**, The encoder sees one cochleogram and outputs, per range cell, spherical-harmonic coefficients of the EnergyScape and of four color maps, which are evaluated on the icosphere and blurred. **b**, The measured self-motion fixes the pose of the head at each later call, and the EnergyScape is moved into that pose by an exact warp without parameters. **c**, A renderer with prescribed spreading, learned absorption, range blur and echo spectra, and a learned ERTF (a small network on spherical-harmonic features of the direction) turns the moved EnergyScape into the levels expected at call *t* + *k*. **d**, The squared error with respect to the levels actually heard, together with a sparsity term and two smoothness terms, updates all learned parts (filled boxes). Layer sizes and parameter counts are those of the runs reported here.

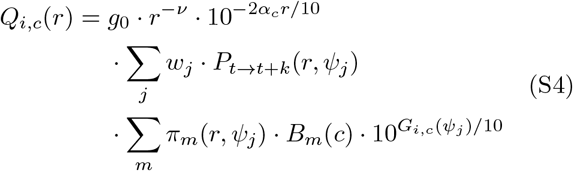

The four echo spectra *B*_*m*_(*c*) are learned (256 parameters, stored in dB with zero mean over bands, initialized with a standard deviation of 1 dB). The spreading exponent *ν* is that of the data (*ν* = 2 here), the one-way absorption *α*_*c*_ per band is learned (64 parameters, initialized at 0.035 dB/m per kHz), and the gain *g*_0_ is fixed and calibrated from the cochleograms alone (Methods). A learned point-spread function *h*_*c*_(*r*) of 13 taps per band (832 parameters, a softmax over taps, initialized with 63% of its weight on the central tap) is then convolved along range, and the result is converted into levels in the same units as the input:

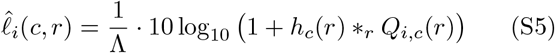

which is equation (6) of the Methods.

#### Loss and Training

Each training example is a window of 17 consecutive calls in one world. The encoder sees only its first call, and the loss compares the levels predicted for calls *t* + *k* with the levels heard, over the range cells beyond 1.06 m (Methods, equation (9)). A sparsity term on *P* ^*raw*^ keeps the scene compact, and two smoothness terms on the ERTF, one across directions and one across bands, penalize steep edges only linearly, so that spectral notches can form. The gradients of this loss update all learned parts together. In the online experiments, one AdamW step is taken after every call, on 16 windows drawn from the replay memory (*N* = 64, the windows that end within the last 65 calls, Methods, equation (10)).

### 11.2 Supplementary Note 2: The Experiments

Unless stated otherwise, all experiments use the point worlds, the smooth paths and the model of the Methods. They learn online with a replay memory of 64 calls over one drive of 60,000 calls through the 40 training worlds, and they are scored on the four unseen validation worlds. Supplementary Table 1 lists the scores of every run.

- *Main online run (E1)*: the agent learns online with the full head motion (pan, tilt and roll), starting from the weak prior. After the first drive, learning continues for a second drive of 60,000 calls through the same worlds with new paths, head motion and noise (120,000 calls in total).
- *Batch reference (E2)*: the same model is trained in batch mode on all windows of the first drive of E1, in shuffled epochs. It needs the whole drive in memory and serves only as a reference for the online learner.
- *Relearning after a change of the ears (E3)*: starting from the end of the 120,000-call drive of E1, the left ear is rotated upward and the right ear downward by *±*5°, *±*10° or *±*20° about the interaural axis, with the emitter unchanged. Online learning continues for 30,000 calls. The ERTF is scored against the rotated and the original ears.
- *Replay memory (E4)*: the first drive of E1 with a memory of *N ∈ {* 1, 4, 16, 256, 512*}* calls (*N* = 1: every step uses only the two newest windows).
- *Seeds (E5)*: the first drive of E1 repeated with network seeds 1 and 2 (E1 is seed 0).
- *Ablations (E6)*: the first drive of E1 without the weak prior (random start) and without the two smoothness terms. The run without the prior was repeated with network seeds 1 and 2.
- *Head axes (E7)*: the first drive of E1 with only some of the head rotations: pan only, pan and tilt, and pan and roll (pan, tilt and roll is E1). The body paths, worlds and noise are those of E1.
- *Symmetric worlds (E8)*: as E1, but in point worlds whose reflectors are spread symmetrically about the height of the sensor (0.40 *±* 2.6 m), with 1.7 times as many reflectors. In the frame of the head, which tilts up more than down, 68% of the isolated echoes then come from below.
- *Specular surfaces (E9)*: two drives of 60,000 calls through sparse point worlds (0.05 reflectors per m^3^, a sixth of the main density), one without surfaces and one inside a room of specular surfaces (a floor, a ceiling at 3.2 m and four walls 3 m beyond the arena) with exactly the same reflectors (Supplementary Fig. 7). Each run is scored on the validation worlds of its own kind.

The five controls are repetitions of the first drive of E1:

- *Shuffled motion (C1)*: the self-motion readings of every window come from a randomly chosen other window of the drive.
- *No prediction (C2)*: only the reconstruction of the current call is learned (ℒ_0_), without the prediction of later calls.
- *Shifted motion (C3)*: every window receives the true self-motion of the window two calls earlier.
- *Head fixed (C4)*: no pan, tilt or roll, on the same body paths.
- *Causal calibration (C5)*: the level scale and the renderer gain are calibrated on the first 1,500 calls only.

**Supplementary Table 1:** Scores of All Runs. End-of-drive scores of the final model of every run (E2: the checkpoint with the lowest validation prediction error). Correlation with the true ERTF in the echo field (*ρ*_EF_), over the whole frontal hemisphere (*ρ*_H_), of the ILD (*ρ*_ILD_) and of the up/down part (*ρ*_UD_) (E3: against the rotated ears, whose table is undefined at the edge of the frontal hemisphere, so no whole-hemisphere score is given). Localization of isolated echoes in the unseen validation worlds from a single call: median great-circle, azimuth and elevation errors (degrees), and the percentage of correct left/right and up/down decisions, next to the percentage obtained by always choosing the more frequent side in elevation. The second drive of E1 is scored after 120,000 calls.

| Run | Correlation with true ERTF | | | | Median error ($^{\circ}$) | | | Correct (%) | | |
| --- | --- | --- | --- | --- | --- | --- | --- | --- | --- | --- |
| | $\rho_{\text{EF}}$ | $\rho_{\text{H}}$ | $\rho_{\text{ILD}}$ | $\rho_{\text{UD}}$ | Bearing | Az. | El. | L/R | U/D | Majority |
| E1 Main run, seed 0 | 0.97 | 0.87 | 0.95 | 0.77 | 4.4 | 2.5 | 2.8 | 95 | 88 | 59 |
| E5 Main run, seed 1 | 0.96 | 0.86 | 0.95 | 0.75 | 3.9 | 2.3 | 2.5 | 95 | 93 | 59 |
| E5 Main run, seed 2 | 0.97 | 0.89 | 0.95 | 0.81 | 4.1 | 2.5 | 2.4 | 96 | 94 | 59 |
| E1 Second drive | 0.97 | 0.87 | 0.95 | 0.80 | 6.0 | 3.3 | 4.5 | 87 | 74 | 60 |
| E2 Batch reference | 0.97 | 0.89 | 0.95 | 0.82 | 2.9 | 1.7 | 1.9 | 98 | 97 | 59 |
| E4 Memory $N = 1$ | 0.91 | 0.85 | 0.91 | 0.66 | 6.4 | 3.4 | 4.1 | 95 | 94 | 59 |
| E4 Memory $N = 4$ | 0.93 | 0.84 | 0.92 | 0.68 | 6.2 | 3.4 | 3.9 | 97 | 92 | 59 |
| E4 Memory $N = 16$ | 0.96 | 0.84 | 0.94 | 0.74 | 5.2 | 2.8 | 3.3 | 96 | 93 | 59 |
| E4 Memory $N = 256$ | 0.97 | 0.86 | 0.95 | 0.78 | 4.4 | 2.8 | 2.9 | 91 | 81 | 59 |
| E4 Memory $N = 512$ | 0.97 | 0.90 | 0.95 | 0.80 | 4.0 | 2.2 | 2.6 | 97 | 96 | 59 |
| E6 No prior, seed 0 | 0.97 | 0.86 | 0.95 | 0.79 | 4.1 | 2.4 | 2.6 | 97 | 93 | 59 |
| E6 No prior, seed 1 | 0.96 | 0.87 | 0.95 | 0.73 | 3.9 | 2.3 | 2.4 | 96 | 94 | 59 |
| E6 No prior, seed 2 | 0.96 | 0.88 | 0.94 | 0.72 | 4.4 | 2.5 | 2.7 | 97 | 89 | 59 |
| E6 No smoothness | 0.91 | 0.77 | 0.93 | 0.63 | 4.4 | 2.5 | 2.7 | 96 | 92 | 59 |
| E7 Pan only | 0.48 | 0.69 | 0.79 | 0.03 | 15.3 | 2.4 | 14.7 | 98 | 44 | 82 |
| E7 Pan and tilt | 0.96 | 0.87 | 0.95 | 0.73 | 4.1 | 2.5 | 2.5 | 98 | 91 | 61 |
| E7 Pan and roll | 0.71 | 0.82 | 0.77 | 0.06 | 5.6 | 2.4 | 4.3 | 97 | 73 | 79 |
| E8 Symmetric worlds | 0.97 | 0.93 | 0.97 | 0.95 | 4.3 | 2.4 | 2.9 | 97 | 94 | 67 |
| E9 Sparse points | 0.90 | 0.77 | 0.90 | 0.54 | 6.7 | 4.0 | 4.4 | 89 | 77 | 68 |
| E9 Sparse points in a room | 0.86 | 0.84 | 0.89 | 0.68 | 8.2 | 4.5 | 5.5 | 93 | 82 | 66 |
| C1 Shuffled motion | 0.22 | 0.51 | -0.08 | 0.16 | 27.8 | 12.4 | 23.2 | 50 | 40 | 59 |
| C2 No prediction | 0.02 | 0.01 | -0.02 | 0.02 | 47.3 | 26.1 | 27.4 | 38 | 37 | 59 |
| C3 Shifted motion | 0.87 | 0.89 | 0.90 | 0.64 | 19.3 | 9.5 | 15.3 | 90 | 57 | 59 |
| C4 Head fixed | 0.51 | 0.63 | 0.80 | -0.02 | 22.9 | 5.6 | 20.8 | 72 | 35 | 82 |
| C5 Causal calibration | 0.97 | 0.87 | 0.95 | 0.78 | 4.2 | 2.3 | 2.7 | 97 | 93 | 59 |
| E3 Ears $\pm 5^{\circ}$ | 0.97 | - | 0.95 | 0.83 | 5.6 | 3.5 | 3.8 | 87 | 77 | 62 |
| E3 Ears $\pm 10^{\circ}$ | 0.97 | - | 0.94 | 0.88 | 5.8 | 3.8 | 3.9 | 88 | 77 | 62 |
| E3 Ears $\pm 20^{\circ}$ | 0.97 | - | 0.93 | 0.94 | 6.2 | 4.3 | 4.0 | 83 | 77 | 65 |

### 11.3 Supplementary Note 3: Simulation and Evaluation Details

Details of the Methods, in the order in which they are used there.

#### Reflectors

Reflector *n* has a backscatter strength 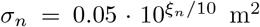 m^2^ with *ξ*_*n*_ normally distributed (standard deviation 5 dB), a size *a*_*n*_ drawn uniformly between 1 and 10 cm that sets a Rayleigh roll-off (*ka*_*n*_)^2^*/*(1 + (*ka*_*n*_)^2^) of the echo amplitude (with *k* the wavenumber), and a random spectral tilt (standard deviation 2 dB between 30 and 90 kHz). Together, roll-off and tilt form *g*_*n*_(*ω*) in equation (1).

#### Received Signals

The signals *s*_*M*_ [*k*] follow from equation (1) by an inverse DFT (*f*_*s*_ = 500 kHz, 32,768 samples), after which white Gaussian noise (standard deviation 2 *·* 10^*−*6^ in the simulator’s amplitude units) is added.

#### Head Movements

Pan and tilt follow sums of two sinusoids with random periods and phases (25–120 calls for pan and 6–30 calls for tilt, overdriven so that the tilt dwells at its limits), tracked at up to 10° per call and interrupted, with a probability of 0.05 per call, by fast moves (6–10° per call) to random targets. The roll is an independent sum of two sinusoids (periods of 10–60 calls) limited to 6° per call.

#### Noise Floor

The noise floor *F*_*i*_(*c*) of ear *i* and band *c* (equation (4)) is the lower tercile over calls of the median power in the 30 farthest range cells, converted from median to mean assuming an exponential distribution.

#### Gain

The fixed gain *g*_0_ of the renderer (equation (6)) is calibrated from the cochleograms alone: a reflector with *P* = 1 over 0.025 sr at an ERTF gain of 0 dB should reproduce the 99.9th percentile of the range-compensated power above the noise floor, 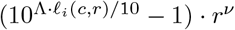, between 0.5 and 2 m.

#### Batch Reference (E2)

The batch run was stopped at step 25,000 of a 60,000-step cosine schedule, resumed under a 120,000-step schedule and stopped at step 64,000 for reasons of compute, when its ground-truth-free validation prediction error (every 1,000 steps, 96 fixed windows of the validation worlds) was no longer clearly decreasing. We report its checkpoint with the lowest validation prediction error (step 63,000).

#### Isolated Echoes

Listed echo contributions within *±*3 cm and 10° are merged into one echo, which counts as isolated if no other contribution within *±*9 cm carries more than 10% of its power, if it lies in front of the head between 0.4 and 7.3 m and if its observed level is at least 10 dB above the noise floor (at most three per call).

#### Scene Evaluation

For the implicit three-dimensional reconstruction, the gain-weighted EnergyScapes of every second call of a validation route were placed in the world frame (with the true poses, for scoring only), keeping per call the cells beyond the receiver blanking (nearer cells are never constrained by the data) that exceed 10% of the call’s maximum. They were accumulated in 10 cm voxels and smoothed by one voxel. Local maxima (5 *×* 5 *×* 5 voxels) above 4% of the 99.9th percentile are detections, matched greedily to the nearest reflector within 0.30 m among those listed as echo sources (1–7.5 m) in at least five calls of the route.

#### Specular Energy

The share of the echo energy carried by the specular surfaces in E9 is one minus the ratio of the cochleogram power without and with the room, for the same routes and reflectors, per band and pooled over the training worlds.

### 11.4 Supplementary Video

#### Supplementary Video 1

The agent learns its ERTF while driving (https://youtu.be/6nxHsedi9fE, also in the Zenodo record). The video explains acoustic flow in a simple scene, shows what the agent infers from single calls along an unseen route (the heard cochleogram, the EnergyScape seen from the head with its acoustic flow and placed in the world, and the predicted against the heard cochleogram), and then plays the learning drive, the main results, the relearning after a change of the ears, the three-dimensional scene the agent found and the effect of specular surfaces. Ground truth is shown for reference only.

### 11.5 Supplementary Figures

**Supplementary Fig. 2:**
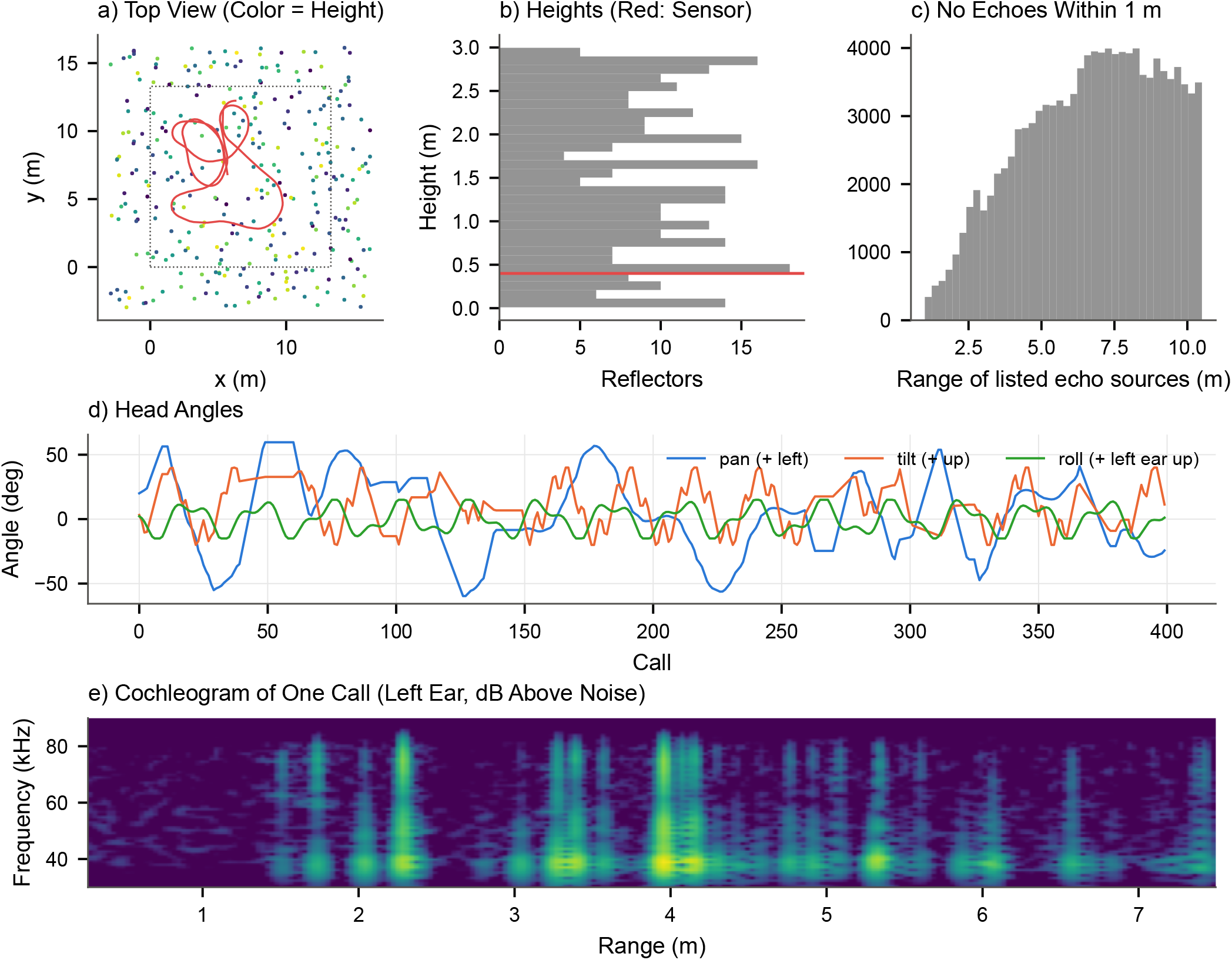
The Simulated Worlds, Paths and Head Movements. **a**, One of the unseen validation worlds from above: point reflectors (color: height) and the first 600 calls of the agent’s smooth path (red). Dotted: the arena. **b**, Heights of the reflectors. Red line: height of the sensor. **c**, Ranges of the echo sources listed by the simulator: no echoes arrive from within 1 m (receiver blanking). **d**, Pan, tilt and roll of the head during the first 400 calls. **e**, The cochleogram of one call (left ear, level above the noise floor).

**Supplementary Fig. 3:**
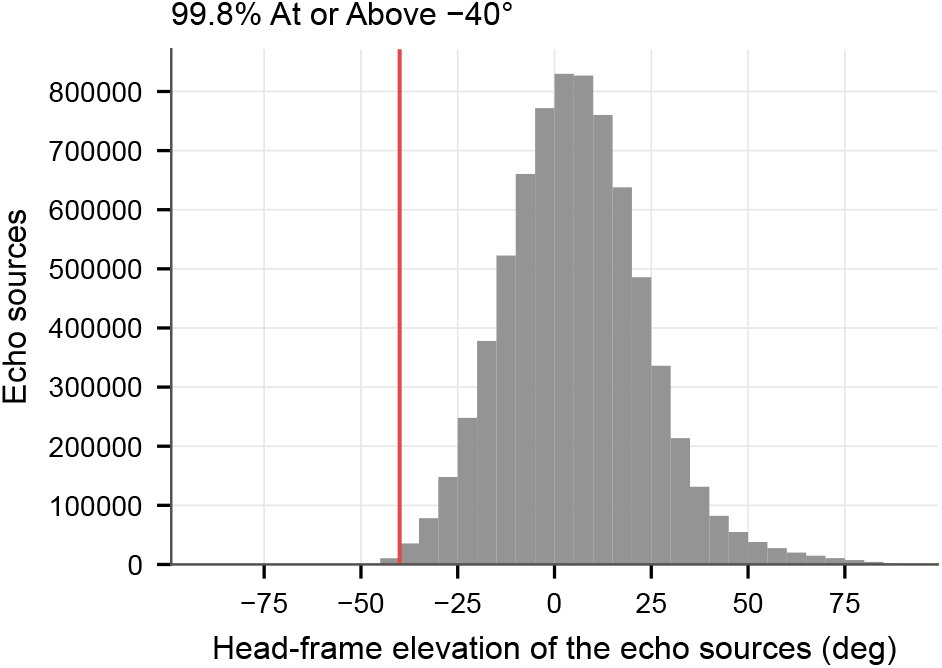
Where the Echoes Come From, and the Echo Field. Head-frame elevation of all echo sources listed by the simulator during the first training drive. Red line: *−*40°. Of the echo sources, 99.8% lie at or above it, because the reflectors stand between the floor and 3 m and the sensor is 0.40 m above the floor. We therefore score the learned ERTF in the echo field: the directions at or above *−*40° elevation and within 25 dB of the maximum gain of both ears (455 directions, 35% of the frontal hemisphere). The echo field is defined from the simulation and used for evaluation only. The learner never sees it.

**Supplementary Fig. 4:**
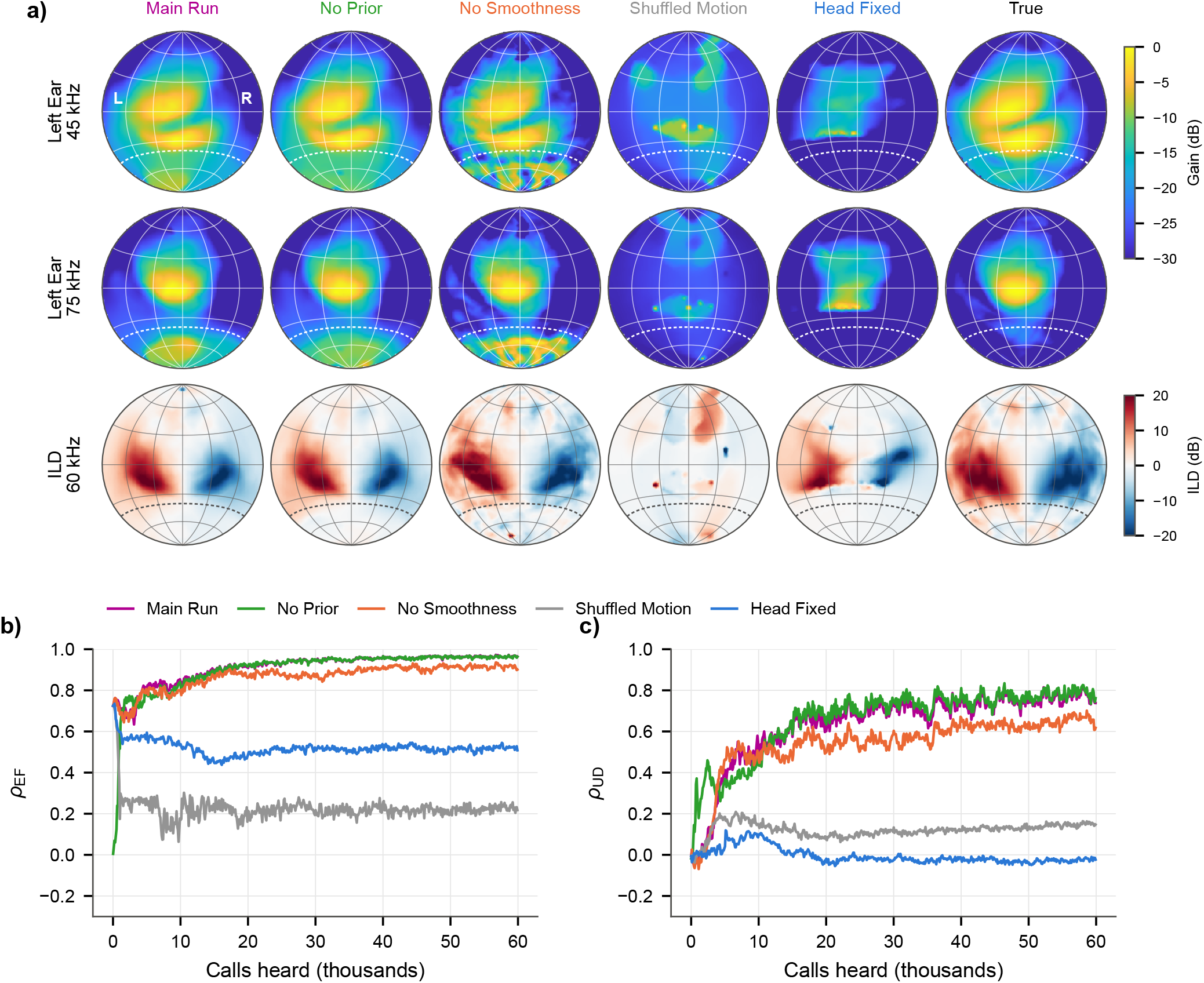
Ablations and Controls. **a**, The learned ERTF after the first drive of 60,000 calls (final model, left ear at 45 and 75 kHz, ILD at 60 kHz) for the main run, without the weak prior (random start), without the smoothness terms, with the self-motion readings taken from a random other moment of the drive, and with the head kept fixed, next to the true ERTF (reference only). Frontal hemisphere seen from behind the head, grid every 30°. Dashed: *−*40° elevation. **b**, Correlation with the true ERTF in the echo field (*ρ*_EF_) over the drive. **c**, The same for the up/down part (*ρ*_UD_). One seed each.

**Supplementary Fig. 5:**
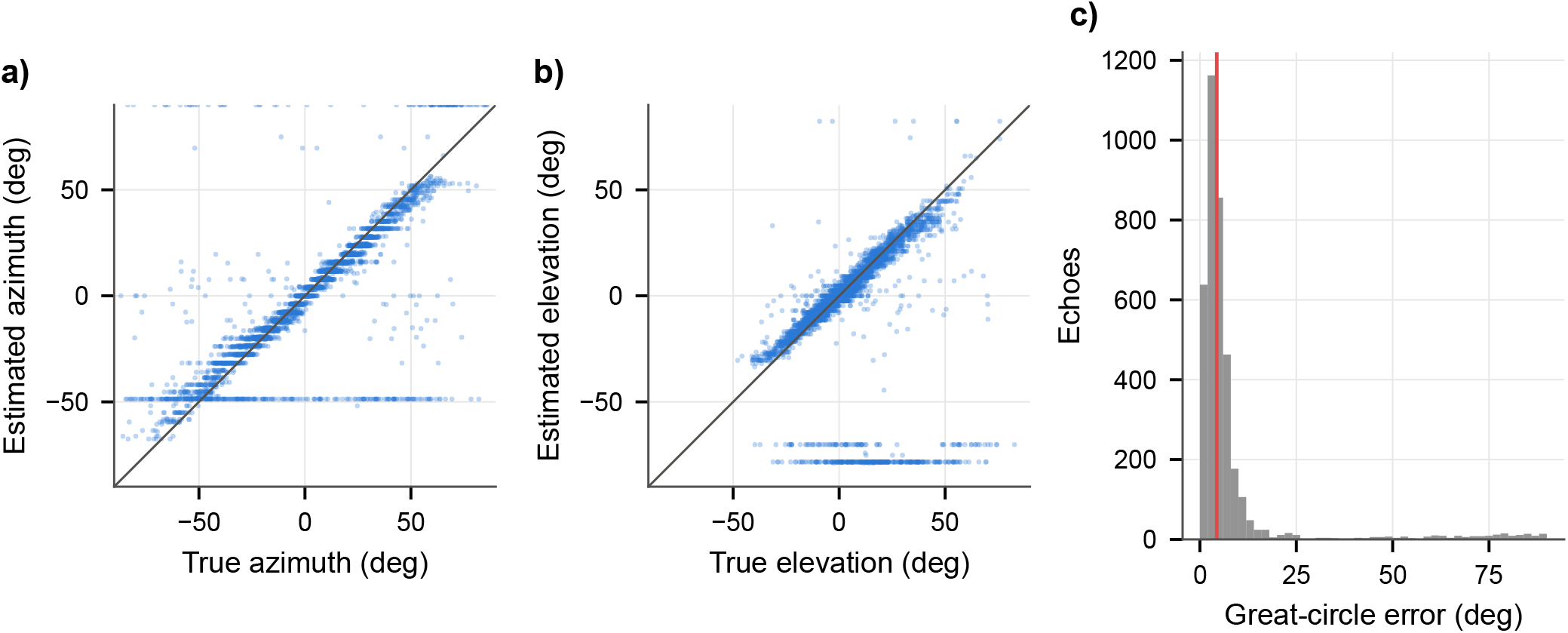
Directions of Isolated Echoes From a Single Call. True against estimated azimuth and elevation of isolated echoes in the four unseen validation worlds after the first drive of 60,000 calls (main run, seed 0), and the distribution of the great-circle error (red line: median).

**Supplementary Fig. 6:**
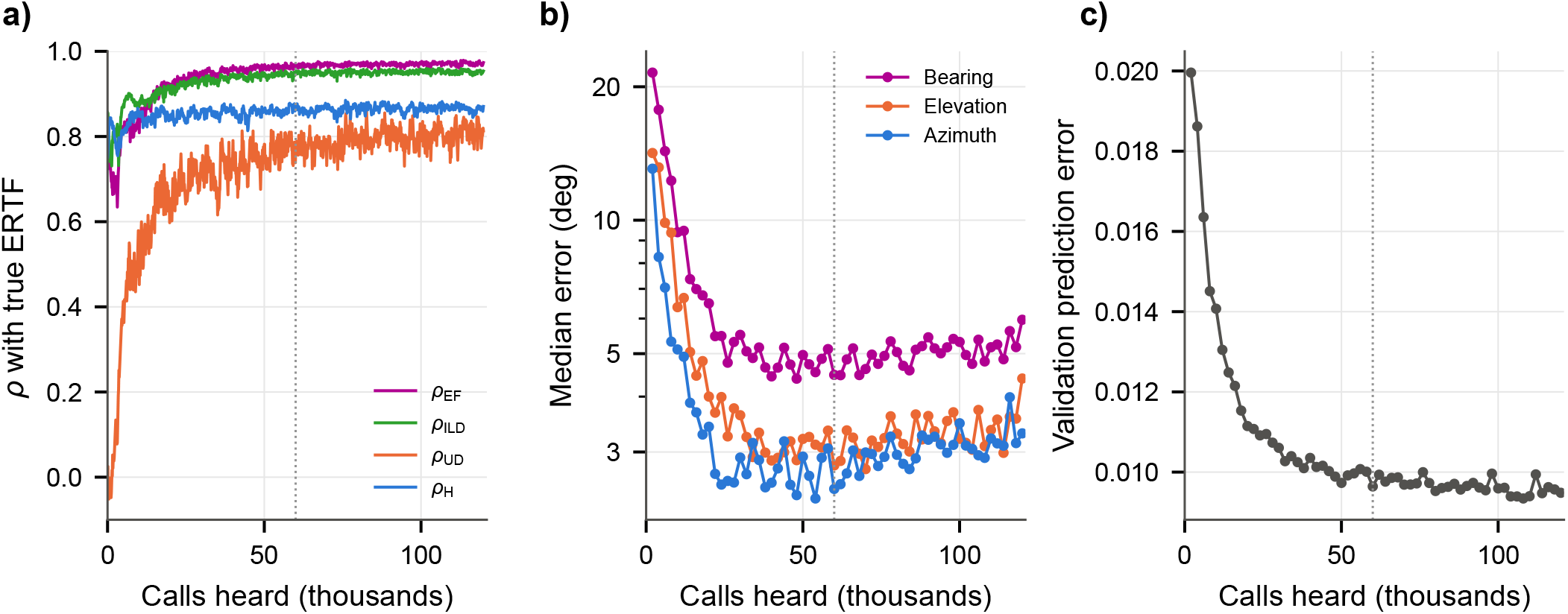
The Main Run Over Two Drives. The online learner of the main run continues for a second drive of 60,000 calls through the same worlds, with new paths, head motion and noise (dotted line: start of the second drive). **a**, Correlation between learned and true ERTF in the echo field (*ρ*_EF_), for the ILD (*ρ*_ILD_), the up/down part (*ρ*_UD_) and over the whole frontal hemisphere (*ρ*_H_). **b**, Median error of the direction of isolated echoes in the unseen validation worlds, inferred from a single call. **c**, The ground-truth-free prediction error on fixed windows of the validation worlds.

**Supplementary Fig. 7:**
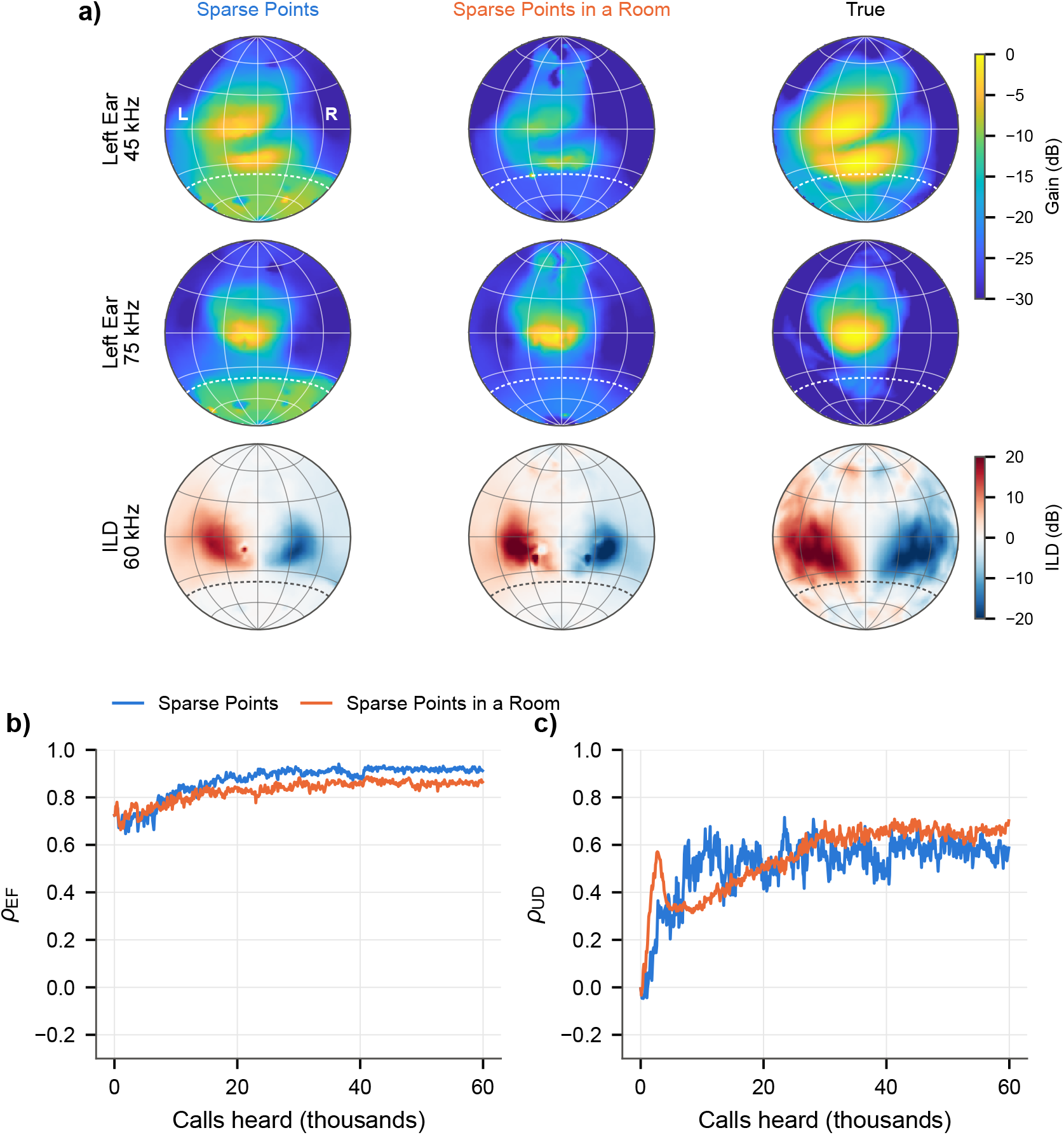
Specular Surfaces Distort the Learned ERTF. **a**, The ERTF learned online in sparse point worlds (a sixth of the main reflector density) without surfaces (left) and inside a room of specular surfaces with exactly the same reflectors (middle), after 60,000 calls (final model), next to the true ERTF (right): left ear at 45 and 75 kHz and the ILD at 60 kHz, frontal hemisphere seen from behind the head, grid every 30°. Dashed: *−*40° elevation. The surfaces carry 37% of the echo energy between 30 and 45 kHz and 9% between 75 and 90 kHz. In the room, the main lobe of the left ear at 45 kHz is largely lost. **b**, Correlation with the true ERTF in the echo field (*ρ*_EF_) over the drive. **c**, The same for the up/down part (*ρ*_UD_).

**Supplementary Fig. 8:**
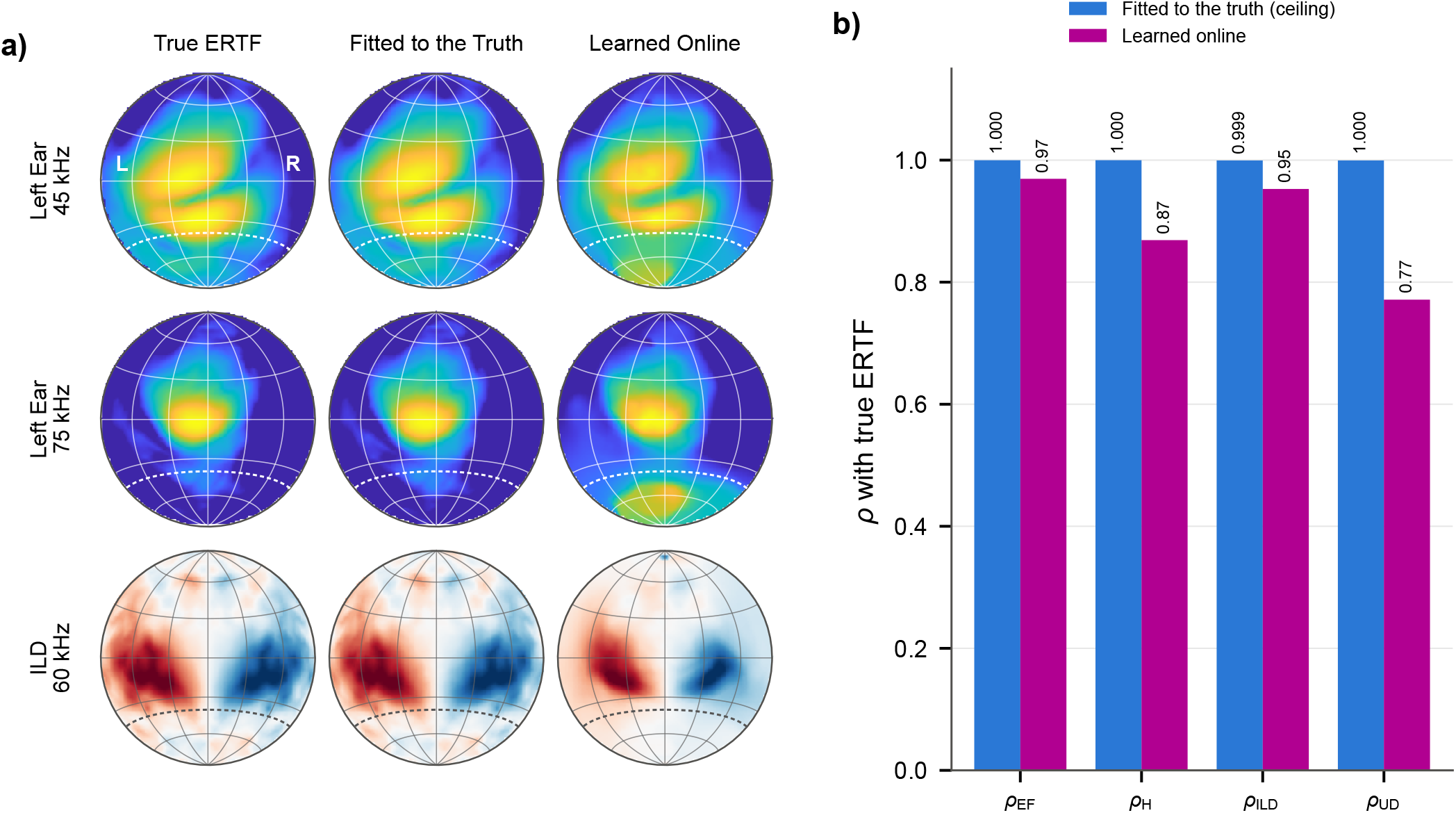
The ERTF Network Can Represent the True ERTF. **a**, The true ERTF (left), the ERTF network of the paper fitted directly to it with the true ERTF as a supervised target (middle, 20,000 Adam steps without the smoothness terms, used only as a check), and the ERTF the main run learned online from its echoes and self-motion after 60,000 calls (right, final model): left ear at 45 and 75 kHz and the ILD at 60 kHz, frontal hemisphere seen from behind the head, grid every 30°. Dashed: *−*40° elevation. **b**, Correlation with the true ERTF in the echo field (*ρ*_EF_), over the whole frontal hemisphere (*ρ*_H_), for the ILD (*ρ*_ILD_) and for the up/down part (*ρ*_UD_). The fitted network reaches 0.999 or more on every measure (0.18 dB RMS), so the differences between the learned and the true ERTF come from the learning, not from the representation.

